# Tumor-derived FXII engages the intrinsic coagulation cascade to support breast cancer liver metastasis

**DOI:** 10.64898/2026.06.03.729507

**Authors:** J. Garcia-Lerena, JR. Jhan, N. Talukdar, J. Vusich, M.M. Ortiz, A.J. Schulte, M. Atkins, D. Patel, D. Hollern, M. Quackenbush, B. To, S. Marei, W. Hui, Y. Lu, T. Kiki-Teboum, MJ Flick, B. Chen, J. Luyendyk, E. Andrechek

## Abstract

Metastasis is the leading cause of death in breast cancer, yet the mechanisms controlling organotropism are not well defined. Coagulation has emerged as a biologically relevant contributor to metastatic progression, but mechanisms linking pro-thrombotic phenotypes to organ-specific metastasis remain unresolved, significantly hindering the development of novel treatments. Here, a serial transplantation approach was used to enrich for liver organotropism from a spontaneous mouse mammary tumor model with occasional liver and lymph node metastasis. Comparative transcriptomics between the enriched liver and lymph node metastases revealed strong upregulation of coagulation in liver metastases, due in part to loss of repression of FXII with knockout of the E2F5 transcription factor. *In vitro* clotting assays demonstrated that tumor-derived FXII was sufficient to induce fibrin(ogen) clot formation. Moreover, liver metastatic cells exhibit elevated lipid peroxide levels and impaired lipid droplet formation associated with a pro-coagulant phenotype. Inhibition of coagulation with low molecular weight heparin reduced the presence of circulating tumor cells and suppressed liver metastasis in the mouse model. Human electronic health record data supported the translational relevance of these findings. Together, these data reveal a new mouse model where loss of E2F5 has resulted in tumors with elevated expression of FXII that have a propensity for liver metastasis and illustrates that anti-coagulation dramatically reduces the liver-specific metastasis in breast cancer.

**Highlights:**

E2F5 conditional knockout model develops breast tumors with liver tropism
Liver metastasis hijacks the intrinsic coagulation cascade mediated by tumor-derived FXII
Liver metastatic cells displayed lipid metabolic alterations that contributed to a pro-coagulant phenotype
Low molecular weight heparin blocks liver metastasis and significantly reduces circulating tumor cells

## Introduction

Progression to distant metastasis remains the main cause of breast cancer mortality ^1,2^. Metastatic progression is a complex cascade ranging from intravasation, to survival in circulation, and extravasation ^3–5^ with organotropic preferences shaping colonization of distant organs. Breast cancer primarily metastasizes to the lung, brain, bone and liver ^6^ with cell line experiments revealing mechanisms driving metastasis to brain, bone and lung ^7–9^. Despite the clinical relevance, the mechanisms driving liver metastasis remain poorly understood ^10,11^.

One key limitation is the lack of genetically engineered mouse models (GEMMs) that spontaneously metastasize to the liver ^12–16^. Indeed, most GEMMs used to study metastasis, including MMTV-Neu, MMTV-Wnt-1, and MMTV-PyMT, preferentially develop pulmonary metastasis ^13–17^. While there are models where low levels of liver metastasis have been reported, including the conditional Cdh1/p53 knockout line (KEP) (Derksen et al., 2006) and the p53 mammary-specific knockout ^18^, the majority of spontaneous models with an intact immune system lack liver metastasis, preventing the study of the mechanisms driving liver tropism. Currently, intrasplenic and intraportal injection models are commonly used to study liver colonization and, while valuable, these models have key limitations as they do not recapitulate the complete metastatic cascade ^19^. In these injection approaches, human MDA-MB-231 and mouse 4T1 lines are commonly used but preferentially develop pulmonary metastasis ^20,21^. Additionally, cell injections are often performed on immune-compromised models that lack a physiologically relevant tumor microenvironment, forcing immune transplantation strategies ^12^. This underscores the need for a spontaneous mouse model that develops metastasis to the liver.

Liver metastases from a variety of primary cancers have conserved histopathological growth patterns that are largely defined by the tumor-liver interface ^22^. The main growth patterns are desmoplastic (DGP) and replacement (RGP) ^23^. Mammary splenic injections with the 4T1 cell line have shown RGP with distinct metabolic adaptations that enabled liver metastasis ^24^. Metastatic RGP lesions are characterized by low angiogenesis, co-option of sinusoidal blood vessels, and the replacement of hepatocytes with tumor cells that mimic liver architecture ^25,26^. This growth pattern is observed in >90% of human breast cancer liver metastases ^27^. Vascular co-option by cancer cells induces upregulation of angiopoietin-2 (Ang2) which antagonizes the Tie-2 receptor, leading to vascular damage, apoptosis, and correlates with intravascular thrombosis ^28^.

Thrombosis is associated with the dysregulation of the coagulation cascade, and the liver has a central role in hemostasis by producing most of the coagulation factors involved ^29–32^. The function of the coagulation cascade in shaping the tumor microenvironment is well studied ^33–36^, with recent findings illustrating that aspirin may prevent metastasis by acting through platelets ^37^. Additionally, tumor cell-educated platelets induce macrophage recruitment and secretion of pro-inflammatory cytokines in melanoma ^38^. Notably, active FXII (FXIIa) has been reported to trigger macrophage secretion of TNF-α, IL-6, IL-12, and IL-1β and promote atherosclerosis in mice ^39^. Moreover, human monocytic cells exhibited transformation to M2-like phenotypes (IL-4^high^, IL-10^high^, transforming growth factor beta (TGFβ)^high^, TNF-α^high^) when stimulated with coagulation factor VIIa (FVIIa), FIII, or FXIIa ^40^.

Despite these advances, the mechanisms describing drivers of breast cancer liver metastasis are limited, in part by the lack of spontaneous mouse models with liver tropism. This study reports a new mouse model of spontaneous breast cancer liver metastasis and identifies the role of the intrinsic coagulation pathway as an organotropic mechanism, contributing to the migration, survival and invasion of breast cancer cells to the liver.

## Materials and Methods

### Animal studies

Tumors were orthotopically implanted into the abdominal mammary gland of 8- to 12-week-old MMTV-Cre female mice. Mice were palpated twice a week for mammary tumor formation, and weight loss and behavior were monitored. Tumor sizes were measured weekly using calipers along two perpendicular axes: length (the longest dimension) and width (the shorter dimension). When mammary tumors reached 2000 mm^3^, samples were harvested and stored in DMEM with 20% FBS and 10% DMSO at −80 °C before long-term storage in liquid nitrogen vapor phase, or flash frozen in liquid nitrogen for DNA, RNA or protein extraction. Animals were euthanized using CO_2_ inhalation followed by cervical dislocation as a secondary method except when blood was collected, and necropsy was immediately performed. Next, tumors were formalin-fixed and embedded in paraffin for histological analysis. All animal studies were approved by the Institutional Animal Care and Use Committee (IACUC) under protocol number PROTO202400120.

### Cardiac puncture

Mice were humanely euthanized using CO_2_. Immediately after animals stopped breathing, they were removed from the induction chamber and placed in dorsal recumbency, and the thoracic cavity was opened carefully. A 23 G needle attached to a 1 mL syringe was inserted into the heart at a 40° angle to the body and blood was collected and placed in lithium heparin blood collection tubes (Cat. 22-040-021). When plasma was needed, syringes were additionally pre-loaded with 0.3 mL of 3.8% sodium citrate to prevent coagulation. Blood fractionation and colony formation assay

In a 10 mL tube, 1-1.5 mL of anticoagulant-treated blood was added and mixed with an equal volume of balanced salt solution. Diluted blood was carefully layered onto a 10 mL tube containing 3 mL of Ficoll-paque premium medium (Cat. GE17-5446-02). Samples were centrifuged for 40 minutes at 400 g and 18 °C. The upper plasma layer, a second layer containing mononuclear cells, and the bottom layer of erythrocytes were collected for further analysis. For colony formation assays, the mononuclear fraction was washed by adding an equal volume of sterile 1x PBS and centrifuging at 400 g for 15 minutes at 18 °C. The supernatant was discarded, and the cell pellet was resuspended in complete DMEM medium and plated in 10 cm plates for 21 days, changing the medium every 3 days. At 21 days, the medium was aspirated, and plates were washed with 1x PBS. The plates were then incubated with 5 mL of 10% formalin for 25 minutes. The formalin was then removed, and methanol was added for 2 minutes. Plates were incubated for 25 minutes with 6 mL of 0.5% crystal violet solution to stain the circulating tumor cells (CTCs). The staining solution was removed, and plates were washed with tap water. Colonies were visualized under the microscope and quantified. Additionally, DNA was extracted from CTCs with QIAamp DNA FFPE Tissue Kit (Cat. 56404) and Endpoint PCR was performed using the E2F5 excision-specific primers E2F5CKO (F: 5’-TGCCTCCCTTATAATTTTGGCC-3’, R: 5’-ACAGTCAGAGCGCAGACCAGG-3’ and un-excised control F: 5’-TGCCTCCCTTATAATTTTGGCC-3’, R: 5’-AGAATCTCATCAAAAGGCAGCC-3’ pair) and Platinum Superfi II Green PCR Master Mix (Thermo Fisher Cat. 12369010) to determine the E2F5 CKO origin of CTC colonies.Cell culture and E2F5 CKO primary cell line generation

### Cell Culture

E2F5 CKO primary cells were cultured in DMEM low glucose (Sigma Cat. FG0415), 10% fetal bovine serum (Corning, Cat. 35-015-CV), and 1% Penicillin-Streptomycin (Life Technologies, Cat. 15240062). MCF-7 and MDA-MB-231 human cell lines were cultured in RPMI medium (Corning #50-020-PC), 10% fetal bovine serum, 1% Penicillin-Streptomycin, 23.8 mM sodium bicarbonate, 25 mM HEPES, and 1 mM sodium pyruvate. Cells were incubated at 37 °C and 5% CO_2_. For primary cell generation, E2F5 CKO tumor pieces of 2 mm^3^ were cut with sterile surgical tools and placed in a 10 cm culture dish with 5 mL of complete DMEM low glucose (Sigma Cat. FG0415) to allow cells to dissociate. The time of dissociation varied between 4-7 days depending on the tumors. Cells were passaged several times to enrich the epithelial cell population by applying differential trypsinization using 1.5 mL of 0.05% trypsin and incubating 5 minutes at room temperature. Trypsin sensitive subpopulation was removed, likely containing fibroblasts and stromal cells. Plates were washed with PBS and fresh medium was added. Cells were allowed to recover for two days, and the process was repeated three times. Next, DNA from the trypsin-resistant population was extracted with Quick-DNA Miniprep Plus Kit (Zymo Research Cat. D4069) and Endpoint PCR was performed using the E2F5 excision-specific primer E2F5CKO, (F: 5’-TGCCTCCCTTATAATTTTGGCC-3’, R: 5’-ACAGTCAGAGCGCAGACCAGG-3’ and un-excised control F:5’-TGCCTCCCTTATAATTTTGGCC-3’, R: 5’ AGAATCTCATCAAAAGGCAGCC 3’ pair) and Platinum Superfi II Green PCR Master Mix (Thermo Fisher Cat. 12369010) to determine the E2F5 CKO origin. Sanger sequencing was used to corroborate loss of E2F5 in isolated cell lines. Tumor cell lines were further validated for tumorigenic potential by mammary fat pad injections into MMTV-Cre recipient mice, in which they formed tumors.H&E Staining

### Pathology

Formalin-fixed tissues embedded in paraffin were sectioned at 4-5 µm and mounted on charged slides. Sections were dried in a 56 °C slide incubator for 2 – 24 h and stained on the Leica Autostainer XL using the Hematoxylin and Eosin staining method. For this, slides were incubated twice for 5 minutes with xylene, followed by two incubations with absolute ethanol for 5 minutes each, 95% ethanol for 2 minutes, and rinsed with tap water. Slides were stained with CATHE Hematoxylin 1:10 (BioCare) for 1.5 minutes, followed by a 10-second differentiation in 1% aqueous glacial acetic acid and tap water for 2 minutes. The slides were then covered in 95% ethanol for 2 minutes, stained with 1% Alcoholic Eosin-Phloxine B for 2 minutes, followed by one change of 95% ethanol for 2 minutes, 100% ethanol four times for 2 minutes each, and four changes of Xylene for 2 minutes each before coverslipping with mounting medium (Fisher Scientific, Cat. SP15-500). Slides were scanned using the Olympus SLIDEVIEW VS200 at 40x magnification and brightfield.

### Immunohistochemistry

The slides mounted with paraffin sections were deparaffinized and rehydrated with decreasing ethanol gradients. Next, samples were incubated with 3% hydrogen peroxide in methanol for 10 minutes at room temperature to block endogenous peroxidases. Afterward, slides were rinsed with PBS twice for 5 minutes. Different pre-treatment methods were used for retrieval during optimization: no pretreatment, proteolytic enzyme retrieval, and high-heat retrieval in a vegetable steamer or microprocessor-controlled pressure cooker at pH 6.0 and pH 9.0 depending on the antibody. Samples were blocked with 5% BSA for 30 minutes at room temperature. Next, slides were incubated with primary antibodies against Vimentin (1:1800, Abcam Cat. ab92547), CD31 (1:50, American Research Products Cat. DIA-310), Ki67 (SP9) (1:250, Sigma Cat. 275R-15), CK19 (1:600, Abcam Cat. ab52625), Fibrinogen (1:400, Agilent Technologies Cat. A0080), Perilipin2/ADFP (PLIN2) (1:250, Novus Biologicals Cat. NB110-40877), F4/80 (1:200, Cell Signaling Cat. 70076), TNFα (1:200, Novus Biologicals Cat. NBP1-19532) and FXIIa (1:300, Proteintech Cat. 12551-1-AP) diluted in 1% BSA in PBS-Tween-20 0.1%. Slides were incubated with avidin-peroxidase complex, and 3,3’-diaminobenzidine substrate solution was applied to sections to develop signal color. Counterstaining was performed with CATHE Hematoxylin 1:10 (BioCare) for 1-2 minutes. Slides were dehydrated through decreasing ethanol concentrations and mounted with Permount mounting medium (Fisher Scientific, Cat. SP15-500). Slides were scanned using the Olympus SLIDEVIEW VS200 at 40x magnification and brightfield. DAB signal quantification was performed on ImageJ/FIJI 1.54p software.

### RNA-sequencing

Flash-frozen tumor pieces were homogenized using Fisher Homogenizer 150 (Thermo Scientific). Samples were kept on dry ice to avoid degradation until the homogenizer was used. Total RNA was isolated using QIAGEN RNeasy Midi kit (Cat. 75144) with the manufacturer’s protocol. E2F5 CKO primary cells were seeded in triplicate on 10 cm plates at a density of 2 × 10^6^ cells/well in complete medium overnight. Next, total RNA was isolated using QIAGEN RNeasy Mini Kit (Cat. 74104) with the manufacturer’s protocol. RNA concentration was measured by Qubit (Thermo Scientific) and Agilent 2100 Bioanalyzer. Library preparation and paired-end RNA sequencing (Illumina PE150) were conducted by Novogene using NovaSeq X Plus Series (Illumina) using a minimum coverage of 30M reads per sample. Raw FASTQ files were processed using the nf-core/rnaseq pipeline (Ewels et al., 2020). Adapter and quality trimming was performed using Trim Galore. Trimmed FASTQ files were aligned to the mouse GRCm39 reference genome using STAR. RNA sequencing transcripts were quantified using Salmon. Differential expression analysis was performed using DESeq2 1.38.1 and differentially expressed genes (DEGs) with padj < 0.05 and log_2_ fold change > 2 or < −2 were considered significant. The R package fgsea 1.24.0 was used for Gene Set Enrichment Analysis on DEGs and MSigDB gene sets, including C2 curated pathways and C6 oncogenic signature. Pathways with padj < 0.05 were considered enriched. Gene clusters were analyzed with topGO 2.50.0 for identification of Gene Ontology pathways. Additionally, Cluster 3.0 was used to perform unsupervised hierarchical clustering. RNA sequencing results from human data were obtained from Gene Expression Omnibus accession number GSE209998 ^41^.

### CUT&RUN and Sequencing Library Preparation

CUT&RUN was performed using the CUTANA ChIC/CUT&RUN kit (Epicypher, version 3), following the user manual version 3.1. Briefly, cells were bound to concanavalin-A magnetic beads, permeabilized with 0.01% digitonin, and 5 × 10^5^ HC11 or MCF-7 cells per target (n=2 per target) were incubated at 4 ° C on a nutator with 0.5 µg of antibody overnight in 50 µL of antibody buffer. Cells were washed and pAG-MNase was added to the cells in 50 µL of permeabilization buffer and incubated for 10 minutes at room temperature. Cells were washed again and CaCl_2_ was added to activate MNase and incubated at 4 °C on a nutator for 2 h. The reaction was stopped with EGTA STOP buffer. E. coli DNA was included in the STOP buffer as a spike-in control. Following purification, DNA was quantified using the Qubit dsDNA HS kit. CUT&RUN sequencing libraries were prepared using the CUTANA CUT&RUN Library Prep kit (Epicypher, version 1), following the user manual version 1.4. Briefly, 5 ng of CUT&RUN DNA per target replicate was added in equal volumes for end repair for 20 minutes at 20 °C. Following adapter ligation for 15 minutes at 20 °C and U-excision for 15 minutes at 37 °C, DNA was purified using SPRIselect magnetic beads (Beckman Coulter). Next, library amplification was performed for 14 cycles of indexing PCR, followed by PCR cleanup using 1x volume of SPRIselect beads. Sequencing libraries were analyzed by TapeStation (Agilent) and sequenced on the NovaSeq 6000 (Illumina) at 2 x 50 bp for 8-10 million reads per library.

### CUT&RUN Analysis

Raw FASTQ files were processed using the nf-core CUT&RUN pipeline (nf-core/cutandrun v3.2.2). Adapter and quality trimming was performed using Trim Galore v0.6.6. Post-trim quality control was performed using FastQC v0.12.1. Trimmed FASTQ reads and spike-in reads were aligned to mm10 and K12-MG1655, respectively, using Bowtie2 v2.4.4. Peaks were called using SEACR v1.3, and consensus peaks were merged using bedtools v2.31.0. Genomic tracks were visualized using IGV desktop v2.16.1.

### Cell-based Fibrin Formation and Fibrinolysis Assay

E2F5 CKO primary cells, and human breast cancer cell lines MCF-7 and MDA-MB-231 were resuspended in complete medium at a concentration of 33,000-50,000 cells per 200 microliters and cells were seeded in 8-well chamber slides Lab-Tek II #1.5 coverglass (Thermo Fisher Cat. 155360). Empty wells were filled with PBS to control temperature. Cells were placed in CO_2_ incubator overnight. Next, medium was removed and cells were washed twice with 1x PBS and incubated with Human pooled plasma (Normal 1:5; (Innovative Research Cat. ISERAB100ML), FXII deficient 1:9; (Siemens Cat. 23-044-686, and Precision Biologic Cat. FDP12-10)) 10 mM of sterile CaCl_2_ and 10 mg/150mL of Alexa Fluor 488-Fibrinogen (Fisher Scientific Cat. F13191) or Alexa Fluor 647-Fibrinogen (Invitrogen Cat. F35200). Cells were incubated in complete DMEM or DMEM medium without glutamine (Gibco A1443001) by adding 1g/L of glucose. Cells were also treated with dabigatran (DAB) (Cayman Chemical Cat. 17133), tissue factor pathway inhibitor (TFPI) (Fisher Scientific Cat. 50-208-7840), corn trypsin inhibitor (CTI) (Santa Cruz Cat. Sc-204358), tumor necrosis factor α (TNF- α) (Thermo Fisher Cat. 315-01A-20UG), and IL1β (PeproTech Cat. 211-11B-10µg). Fibrin formation was monitored at different times (30 minutes, 2, 4, and 24 h). Following incubations, cells were washed twice with 1X PBS, fixed with 4% paraformaldehyde for 30 minutes, and mounted with Prolong Gold Antifade with DAPI (Invitrogen #P36935). Wells were imaged using the inverted Nikon A1 confocal microscope at the cell surface and at multiple depths above cell surface. All image parameters (e.g., laser intensity and gain) were kept the same, and several random fields per replicate were imaged. Fluorescence was quantified with the ImageJ/FIJI 1.54p software.

### qRT-PCR

E2F5 CKO primary cells were seeded in triplicate on 10 cm plates at a density of 0.8 × 10^6^ cells/well in complete medium overnight. Total RNA was isolated using QIAGEN RNeasy Mini Kit (Cat. 74104). Flash-frozen tumor pieces were homogenized using Fisher Homogenizer 150 (Thermo Scientific) and total RNA was isolated using QIAGEN RNeasy Midi (Cat. 75144). RNA extraction was performed according to the manufacturer’s instructions. cDNA generation was performed with QIAGEN OneStep RT-PCR Kit (100) (Cat. 210212) and qPCR was performed with PowerUp SYBR Green Master Mix (Life Technologies Cat. A25742) with cDNA as template and 100 nM of primers: (F12, F: 5’-CATCACCTACCAGCACGACTTG-3’, R: 5’-ACCTCGCAGAGCACTGTCTCAG-3’; E-cadherin, F: 5’ CAAAGTGACGCTGAAGTCCA 3’, R: 5’ ACATGAGCAGCTCTGGGTTG 3’; βActin, F: 5’ CATTGCTGACAGGATGCAGAAGG 3’,R: 5’ TGCTGGAAGGTGGACAGTGAGG 3’) following the manufacturer’s protocol. ΔΔCt analysis was used to determine differences in expression as described by (Livak & Schmittgen, 2001).

### Protein extraction and concentration

For cell lysates, E2F5 CKO primary cells and MCF-7 cell line were washed with ice-cold HEPES-Buffered Saline (HBS), scraped from the dish, and resuspended in ice-cold HBS. The cell suspension was centrifuged at 1,000 x g for 5 minutes at 4 °C, followed by lysis on ice with ice-cold Pierce RIPA buffer (Thermo Scientific #89901) with Halt Protease and Phosphatase inhibitor cocktail (Thermo Scientific #78441) for 20 minutes. Cell extracts were centrifuged at 16,000 x g for 20 minutes at 4 °C, and the supernatant was collected. For culture media, four volumes of cold acetone were added and incubated on ice for 30 minutes. Samples were centrifuged at high speed (> 12000 g) for 15 minutes. The pellet was washed with cold acetone, air dry and resuspended in Pierce RIPA buffer (Thermo Scientific #89901). Samples were further concentrated by using the Pierce™ Protein Concentrators PES, 10K MWCO (Thermo Fhisher Cat. 88517) according to the manufacturer’s instructions. Protein quantification was performed using Pierce BCA Protein Assay (Thermo Scientific #23225) according to the manufacturer’s instructions.

### Immunoblotting and densitometry

Immunoblotting was performed as described ^42^. The following dilutions were used for the primary antibodies: FXIIa 1:1000 (Proteintech # 12551-1-AP), Thrombin 1:1000 (Abcam # ab92621), LDLR 2 µg/mL (Thermo Fisher # PA5-22976), FASN 1:1000 (Thermo Fisher # MA5-14887), Beta-ACTIN 1:1000 (Cell Signaling #4967), Vinculin 1:1000 (Cell Signaling #13901). Bands were detected using fluorescent IRDye® 800CW goat anti-rabbit (LICOR #926-32211) secondary antibody diluted 1:10000 in TBS-Tween 0.1% and 5% BSA. Blots were imaged using the LICOR Odyssey M. For total protein stain, the membrane was dried on filter paper 10 minutes at 37 °C. Next, the membrane was rinsed with methanol and washed with DI water and stained with No-Stain Protein Labeling Reagent (Invitrogen Cat. A44449) 10 minutes on a shaker. The membrane was washed three times with dH_2_O and incubated with LICOR premade blocking buffer for 1 h at room temperature. Antibody incubations and imaging were performed using the same conditions. Densitometry was performed using ImageJ/FIJI 1.54p. The mean gray values for the target proteins were corrected to the endogenous control (β-actin in cell lysates and total protein in conditioned media) for data normalization.

### Capillary western blotting

E2F5 CKO tumors were tested for cross-linked fibrin(ogen) as previously described by ^43^. Samples were resolved using Wes 66-440 kDa 25-capillary gels (Protein Simple, San Jose, CA) and fibrin(ogen) levels were determined using polyclonal fibrinogen antibodies for each fibrinogen chain (Proteintech, Chicago, IL). Fibrinogen α and γ chains were recognized at 1:100 dilution, and the β chain at a dilution of 1:5000. Polyclonal rabbit anti-human fibrinogen antibody (Agilent Technologies) was used as described by ^44^. A goat anti-rabbit HRP-conjugated secondary antibody (1:700 dilution) (Jackson Immunoresearch) and Wes Rabbit Master Kit’s reagents were used according to the manufacturer’s protocol (ProteinSimple).

### Reactive oxygen species quantification

Oxidative stress was detected with CellROX Green reagent (Invitrogen Cat. C10444) according to the manufacturer’s instructions. CellROX dye is weakly fluorescent in reduced states and bright upon oxidative events induced by ROS. Briefly, 5 × 10^4^ E2F5 CKO primary cells were seeded in 8-wells chamber slides Lab-Tek II #1.5 coverglass (Thermo Fisher Cat. 155360) with 200 µL of complete DMEM overnight. Next, the medium was replaced with fresh complete DMEM or DMEM without glutamine and incubated for 24 h. CellROX Green reagent was added to a final concentration of 5 μM for ROS detection, and cells were incubated for 30 minutes at room temperature with 1 μg/mL Hoechst 33342 solution (Invitrogen Cat. R37605). Slides were washed three times with PBS, fixed with 4% paraformaldehyde for 30 minutes, and mounted with ProLong glass antifade mountant (Invitrogen Cat. P36982). Confocal images were taken on a Nikon A1 confocal microscope withing 2 h of the CellROx green reagent incubation and imaging was repeated the next day to check for signal intensity changes. Fluorescence was quantified using ImageJ/FIJI 1.54p software.

### Nutrient limitation studies

E2F5 CKO primary cell lines were seeded at an appropriate density for 70-80% confluence in 96-well plates with DMEM low glucose (Sigma Cat. FG0415), 10% fetal bovine serum (Corning, Cat. 35-015-CV) and 1% Penicillin-Streptomycin (Life Technologies, Cat. 15240062). Cells were transferred to experimental medium (DMEM medium without glucose and glutamine (Gibco A1443001)) and completed with 10% dialyzed fetal bovine serum (Gibco, Cat. 26400-044). Glucose- and glutamine- containing conditions were prepared by adding 1 mM glutamine (Thermo Scientific, Cat. J60573.22) or 5 mM glucose (Sigma, Cat. D9434-2506) accordingly. Cells were incubated for 24 h, and viability was monitored with CellTiter-Fluor™ Cell Viability Assay (Promega Cat. G6082) according to the manufacturer’s instructions before performing functional assays.

### Cellular Glutathione Detection

Cellular glutathione (GSH) levels were quantified using the Cellular Glutathione Detection Kit (Cell Signaling Technology, Cat. 13859) following the manufacturer’s instructions. Briefly, E2F5 CKO primary cells were cultured with complete DMEM medium (Gibco A1443001) supplemented with 1mM glutamine (Thermo Scientific, Cat. J60573.22), 5 mM glucose (Sigma, Cat. D9434-2506), 10% dialyzed fetal bovine serum (Gibco, Cat. 26400-044), and 1% Penicillin-Streptomycin (Life Technologies, Cat. 15240062). For glutamine-deprived conditions, the amino acid was not added. 20,000 cells were seeded in 96-well plates in replicates and incubated for 24 h. Conditional medium was added the next day and incubated 24 h. Next, medium was removed and 50 µL of Digitonin Lysis Buffer was added to each well and plates were incubated on a shaker for 15 minutes. Lysates were clarified by centrifugation at 14,000 rpm for 10 minutes at 4 °C, and supernatants were collected for GSH assay. A standard curve was prepared using GSH standards. For each sample and standard, 50 µL of working solution containing monochlorobimane and glutathione-S-transferase was mixed with 30 µL of lysate in black, clear-bottom 96-well plates, and the reaction was incubated for 60 minutes at room temperature in the dark. Fluorescence was measured using a SpectraMax iD3® plate reader (Molecular Devices) with excitation at 380 nm and emission at 485 nm. Glutathione concentrations were calculated from the standard curve.

### Inorganic Polyphosphate quantification

Inorganic polyphosphate (Poly P) levels in cell and tissue lysates were quantified using the Inorganic Polyphosphate Assay Kit (Abcam, ab284528), which employs a fluorescent dye that forms a complex with Poly P. Kit components, including the Poly P Extraction Buffer, Assay Buffer, fluorescent dye, and Poly P Standard, were equilibrated to room temperature before experiments.E2F5 CKO primary cells were cultured in complete DMEM or glutamine-deprived medium. Cells were collected at 90% confluency using TrypLE (Fisher Scientific, Cat. 12604021) and pellets were weighed. 250 µL of Extraction Buffer was added and samples were homogenized using Fisher Homogenizer 150 (Thermo Scientific). Samples were centrifuged at 10,000 g for 15 minutes at 4 C, and cleared supernatants were collected. A 100 µL aliquot of each sample was treated sequentially with 2 µL each of RNase and DNase and incubated at 37 °C for 60 minutes. 5 µL aliquots were used to perform agarose gel electrophoresis and assess residual RNA and DNA. When residues were not detected, samples were incubated with 2 µL of proteinase K at 37 °C for 20 minutes and heat-inactivated at 85 °C for 10 minutes. Samples were moved to ice before transferring to a 96-well plate. A Poly P standard curve was generated by diluting the reconstituted Polyphosphate standard to prepare wells containing 0–350 pmol of Poly P in 50 µL Assay Buffer. For all samples and standards, 50 µL reaction mix (47 µL Poly P Assay Buffer + 3 µL Poly P dye) was added to 50 µL of treated sample or standard, and the plate was incubated at room temperature for 10 minutes. Fluorescence was measured in endpoint mode on SpectraMax iD3® plate reader (Molecular Devices) (Ex 415 nm/Em 550 nm), and Poly P concentrations were derived from the standard curve after blank subtraction. Final Poly P content was expressed as nmol/mg of cells.

### Free Fatty acids concentration

Cellular free fatty acid (FFA) concentration was measured using the Free Fatty Acid Assay Kit (Abcam, ab65341). Briefly, 4 × 10^5^ (70% confluence) or 2 × 10^4^ (30% confluence) E2F5 CKO primary cells were harvested for each condition. Samples were homogenized in 200 μL chloroform/Triton X-100 (1% Triton X-100 in pure chloroform) while samples were kept on ice. Samples were centrifuged at 16,000 g for 10 minutes at 4 °C to separate phases. The organic phase (lower fraction) was collected and air-dried at 50 °C in the fume hood. The dried lipids were dissolved in 200 μL of Assay Buffer 5 by vortexing for 5 minutes and kept on ice for measurements following the manufacturer’s protocol.

### ELISA-based Lipid Peroxidation (4-HNE)

Lipid peroxidation levels were quantified by measuring protein adducts of 4-hydroxynonenal (4-HNE) using the competitive ELISA-based Lipid Peroxidation (4-HNE) Assay Kit (Abcam, ab238538). Immediately before use, 1x Conjugate Diluent was prepared by diluting the 100x stock in 1x PBS. A 10 µg/mL 4-HNE Conjugate solution was prepared by diluting the 1.0 mg/mL stock in 1x PBS (e.g., 25 µL in 2.475 mL PBS) and mixed thoroughly. Equal volumes of the 10 µg/mL 4-HNE Conjugate and 1x Conjugate Diluent were combined, and 100 µL of this mixture was added to each well of the microplate. The plate was incubated overnight at 4 °C to allow coating. The following day, the coating solution was removed, wells were washed twice with 1x PBS, and excess fluid was blotted on paper towels. Each well was then blocked with 200 µL of Assay Diluent for 1 h at room temperature. After blocking, the plate was stored at 4 °C, and the Assay Diluent was removed immediately before use. Standards were prepared by serial dilution of the 4-HNE-BSA stock. E2F5 CKO primary cell lysates were obtained with Pierce RIPA buffer (Thermo Scientific #89901) and diluted in Assay Diluent to fall within the linear range of the standard curve. 200 µL of Assay Diluent were added to each well of the 4-HNE-conjugate-coated microplate. Subsequently, 50 µL of standard or diluted samples was added to designated wells, and the plate was incubated for 10 minutes at room temperature under gentle shaking. Following incubation, the plate was washed three times with 250 µL of 1x Wash Buffer. 50 µL of diluted anti-4-HNE primary antibody (1x working concentration) were added to each well, and the plate was incubated for 1 h at room temperature, followed by three additional washes. 100 µL of HRP-conjugated secondary antibody (in Assay Diluent) was added and incubated for another hour under the same conditions, and wells were washed again to remove unbound antibody. To develop the signal, 100 µL of Substrate Solution (pre-warmed to room temperature) were added to each well and incubated in the dark at room temperature for 10 minutes, monitoring color development. The enzymatic reaction was terminated by the addition of 100 µL of Stop Solution to each well. The absorbance was measured immediately at 450 nm using a SpectraMax iD3® plate reader (Molecular Devices). A standard curve was constructed using the absorbance values of the 4-HNE-BSA standards, and sample 4-HNE concentrations were calculated accordingly after background subtraction. Results were normalized to the number of cells due to differences in growth rates among cell lines, and 4-HNE levels were expressed as pmol/1 × 10^5^ cells.

### FXII ELISA

Mouse FXII levels were measured using the Mouse FXII (Coagulation Factor XII) ELISA Kit (Cat# EKF58957-96T; Biomatik). Standards were reconstituted and serially diluted in sample dilution buffer to generate a calibration curve spanning 3.125–200 ng/mL. E2F5 CKO cell culture supernatants were diluted in the same buffer to fall within the standard range. 100 µL of each standard or sample was added to duplicate wells of the pre-coated capture antibody microplate and incubated for 90 minutes at 37 °C. Wells were washed three times with 300 µL of 1x wash buffer. Subsequently, 100 µL of biotinylated detection antibody (diluted as per manufacturer’s instructions) were added and incubated for 60 minutes at 37 °C, followed by three washes. 100 µL of SABC (diluted in SABC buffer) was added and incubated for 30 minutes at 37 °C. After five wash cycles, 90 µL of TMB substrate was added and incubated at room temperature for 20 minutes. The reaction was stopped with 50 µL of stop solution, and absorbance was measured at 450 nm using a SpectraMax iD3® plate reader (Molecular Devices). FXII concentrations were determined from the standard curve after background subtraction. Values were expressed in ng/mL.

### Annexin V Kinetics for PS Exposure

Kinetics of phosphatidylserine (PS) externalization during apoptosis were measured using the RealTime-Glo™ Annexin V Apoptosis/Necrosis Assay (Promega; Cat. JA1000/JA1011). E2F5 CKO primary cells were seeded at 3 × 10^5^ (50 µL per well) in white, tissue-culture-treated 96-well plates (Greiner, Cat. 655083) and incubated overnight. Experimental media (with or without glutamine DMEM) were added, and cells were incubated overnight. Detection Reagent was prepared immediately before use by diluting Annexin V-LgBiT, Annexin V-SmBiT, nano-luciferase substrate, CaCl□, and Necrosis Detection Reagent 500-fold each into complete culture medium (2x final concentration) and mixed gently. Cells were treated with apoptosis-inducing compounds as controls. Next, 100 µL of 2x Detection Reagent was added to each well. Plates were incubated at 37 °C, and PS exposure and membrane permeabilization were measured repeatedly in real time using a SpectraMax iD3® multimode plate reader (Molecular Devices). Background signals from no-cell control wells were subtracted. Kinetic profiles were analyzed by plotting relative luminescence and fluorescence over time.

### Flow cytometry

E2F5 CKO primary cells were cultured for 24 h in complete DMEM or glutamine-deprived DMEM. Media was collected and cells were gently detached using TrypLE (Fisher Scientific, Cat. 12604021). Cells were combined with respective media in 15 mL tubes and centrifuged at 500 g for 5 minutes at room temperature. Next, cell pellets were washed twice with cold Cell Staining Buffer (BioLegend Cat. 420201) and resuspended in Annexin V Binding Buffer (BioLegend Cat. 422201) at a concentration of 1 × 10^6^ cells/mL. A 100 µL aliquot of the cell suspension was transferred into a 5 mL polystyrene tube, and 5 µL of Alexa fluor 647 Annexin V (BioLegend Cat. 640912) was added. Samples were vortex and incubated at room temperature for 15 minutes in the dark. After incubation, 200 µL of Annexin V Binding Buffer was added to each tube. Subsequently, 10 µL of PI solution (BioLegen Cat. 421301) was added and samples were incubated for 15 minutes at 4 °C. Samples were analyzed by flow cytometry on an Attune Cytpix (Thermo Fisher) and data were processed with FlowJo v11 software.

### Immune mass cytometry

Formalin-fixed, paraffin-embedded (FFPE) tumor tissue sections were processed for Imaging Mass Cytometry following the manufacturer’s protocol (Standard Biotools). Briefly, slides were baked at 60 °C for 2 h to remove residual paraffin and then dewaxed in fresh xylene for 20 minutes in a fume hood. Sections were rehydrated through a graded ethanol series (100%, 95%, 80%, and 70%) for 5 minutes each, followed by a 5 minutes wash in Maxpar Water. Antigen retrieval was performed by incubating slides in preheated 1x target retrieval solution (pH 9) at 96 °C for 30 minutes. Slides were cooled to approximately 70 °C for 10–15 minutes, washed in Maxpar Water for 10 minutes, and then rinsed in Maxpar PBS for 10 minutes with gentle agitation. Slides were blocked with freshly prepared 3% BSA in Maxpar PBS for 45 minutes at room temperature. The Maxpar OnDemand Mouse Immune Phenotyping IMC Panel Kit (No. 9100003) was used to characterize the tumor immune microenvironment using the following markers: B220 (RA36B2, 176Yb, 1:500) for B cells, CD4 (BLR167J, 159Tb, 1:100) and CD8 (EPR21769, 162Dy, 1:100) for T cells, F4/80 (D2S9R, 156Gd, 1:100) for macrophages, and Ly-6G (1A8, 166Er, 1:250) for neutrophils. Antibodies were diluted in 0.5% BSA in Maxpar PBS. The antibody cocktail (500 µL) was applied to each slide, and slides were incubated overnight at 4 °C. Next, slides were washed twice in 0.2% Triton X-100 in Maxpar PBS and Maxpar PBS for 8 minutes each. Sections were then stained with Cell-ID Intercalator-Ir (diluted to 250 nM in Maxpar PBS) for 30 minutes at room temperature. After intercalator staining, slides were washed twice in Maxpar Water for 5 minutes and air-dried prior to imaging.

### LA-ICP-TOF-MS instrument setup and sample parameters

Slides were loaded into a Bioimage 266-nm laser ablation (LA) system (Elemental Scientific Lasers, Bozeman, MT, USA) fitted with an ultra-fast low dispersion TwoVol3 ablation chamber and a dual concentric injector (DCI3). The laser ablation unit was coupled to an icpTOF S2 (TOFWERK AG, Thun, Switzerland) inductively coupled plasma time-to-flight (ICP-TOF-MS). Instrument performance was optimized daily using NIST SRM612 glass certified reference material (National Institute of Standards and Technology, Gaithersburg, MD, USA). Tuning of the LA-ICP-TOF-MS system included optimization of torch alignment, lens voltages, and nebulizer gas flow to maximize signal intensity for ^140^Ce and ^55^Mn while maintaining low oxide formation based on the ^232^Th^16^O^+^/^232^Th^+^ ratio (< 0.5). Detailed instrument parameters are listed in Supplemental Table 4. Samples were mounted onto the LA sample holder and loaded into the LA system. The system was purged for 5 minutes prior to analysis. Next, ablation patterns were defined on the standards and samples of interest. For gelatin calibration standards, nine parallel line scans were performed across each standard and corresponding blank with the same parameters: 10- or 20-µm circular laser spot sizes at 80 or 100% laser power with a repetition rate of 100 Hz. The interline spacing was set four times greater than the spot size to prevent overlapping. Reference markers were positioned around the tissue specimens prior to analysis. The same laser parameters were applied, with interline distance equal to the laser spot and no overlap between adjacent lines, ensuring complete sampling of the tissue.

### Acquisition and Analysis of LA-ICP-TOF-MS Data

Data acquisition was performed using TofPilot version 1.3.4.0 (TOFWERK AG, Thun, Switzerland). Resulting LA-ICP-TOF-MS data were stored in the open-source hierarchical data format (HDF5), ensuring compatibility with downstream processing workflows. Post-acquisition data processing was performed with Tofware v3.2.0 and implemented as an extension within Igor Pro (Wavemetric Inc., OR, USA). Data processing comprised: (1) correction of temporal drift in mass peak positions through time-dependent mass calibration, (2) characterization of peak shapes, and (3) fitting and subtracting the mass spectral baseline to remove background. The data were further processed with Iolite version 4.8.6 (Elemental Scientific Lasers, Bozeman, MT, USA). Marker quantification and image generation were performed using the R packages EBImage 4.40.1, FNN 1.1.4, and igraph 2.0.3.

### Low molecular weight heparin in vivo treatments

Liver-tropic tumors (2 mm x 2 mm) were orthotopically implanted into the abdominal mammary gland of 8- to 12-week old MMTV-Cre female mice. Enoxaparin (Winthrop, NDC:00955-1003-10) was administered at a dose of 200 IU/kg every 24 h starting on the day of surgery. Enoxaparin stock solution (30 mg/0.3 mL) was stored at room temperature and the working solution was freshly prepared each day prior to use. Individual injection volumes were calculated based on body weight (200 IU/kg), resulting in approximately 200–300 µL of Enoxaparin at 20 IU/mL in sterile saline per mouse with weights ranging from 20–30 g. Treatments continued daily until the designated sample collection time points (days 25, 35, and 38). Mice were palpated twice a week for mammary tumor formation. Weight loss, bleeding complications, and behavior were monitored daily. Tumor sizes were measured every three days using calipers along two perpendicular axes: length (the longest dimension) and width (the shorter dimension). The endpoint was defined as a tumor size equal to 2000 mm on any measured axis for untreated groups, and all animals were euthanized. Ulcerated tumors were discarded. Animals were euthanized using CO_2_ inhalation, and blood was collected by cardiac puncture immediately after CTC identification. To avoid overestimation, tumor volumes were measured ex vivo using the formula volume = 0.5 × length × width^2^. Afterward, tumors and organs were flash-frozen in liquid nitrogen, viable-frozen in complete DMEM medium with 20% FBS and 10% DMSO, or formalin-fixed and paraffin-embedded for histological analysis. All animal studies were approved by the Institutional Animal Care and Use Committee (IACUC) under protocol number PROTO202400120.

### Retrospective cohort study

Retrospective cohort studies were conducted using de-identified electronic health record (EHR) data from the Truveta platform, as previously described ^45,46^. The association between thrombosis and the hazard of developing liver metastasis among patients diagnosed with breast cancer was evaluated. Patients with breast cancer were identified using International Classification of Diseases (ICD) codes corresponding to malignant neoplasm of the breast (ICD-10-CM C50; ICD-9-CM equivalents where applicable) with a first recorded diagnosis between January 1, 2020, and December 31, 2021. Eligible patients were required to be between 18 and 90 years of age at diagnosis and to have a valid recorded gender. Patients with missing race and ethnicity information, no documented follow-up encounters after breast cancer diagnosis, or no available body mass index (BMI) records were excluded. Patients with evidence of liver metastasis or any other malignancy prior to breast cancer diagnosis were also excluded. Thrombosis was defined by the presence of diagnostic codes indicating venous thromboembolism (VTE), including deep vein thrombosis (DVT), and pulmonary embolism (PE), recorded at any time prior to the development of liver metastasis or censoring. The primary outcome was incidence of liver metastasis, identified by the first occurrence of ICD-10-CM code C78.7. Time to liver metastasis was calculated from the date of breast cancer diagnosis to the date of first documented liver metastasis. Patients without liver metastasis were censored at the date of the last recorded clinical encounter. The association between thrombosis and time to liver metastasis was evaluated using Cox proportional hazards for regression models. Survival time was defined as the interval from breast cancer diagnosis to the development of liver metastasis or censoring. Multivariable Cox models were adjusted with baseline covariates which include age at breast cancer diagnosis, ethnicity, BMI (Missing BMI values were imputed using the multivariate imputation by the chained equations (MICE) method), and comorbidity known to influence cancer progression or thrombotic risk, including type 2 diabetes mellitus, hypertension, hyperlipidemia, hepatitis, and chronic liver disease. Cancer-related treatments, including surgery, chemotherapy, and radiation therapy, were also included as covariates. All covariates were assessed prior to the occurrence of liver metastasis. Hazard ratios (HRs) with 95% confidence intervals (CIs) were estimated. Statistical significance was defined as a two-sided p-value <0.05.

For the association of heparin exposure and hazard of liver metastasis among pancreatic cancer patients, subjects aged 45 years or older were identified with a valid ICD-10-CM code for pancreatic cancer (C25) and a first diagnosis date between 2019 and 2021. For the breast cancer cohort, female patients between 18 and 90 years of age at diagnosis were identified using codes for malignant neoplasm of the breast (ICD-10-CM C50; ICD-9-CM equivalents where applicable) with a diagnosis date between January 1, 2020, and December 31, 2021. Patients with evidence of liver metastasis or any other malignancy prior or at the moment of diagnosis were excluded. The index date was defined as the first recorded diagnosis date. The exposure of interest was Enoxaparin NDC:00955-1003-10. The main outcome of interest was the development of LM, identified by the presence of ICD-10-CM code C78.7. Subjects require a minimum of 2 exposure records before LM diagnosis and any time before the last follow-up date. Baseline covariates included age at breast cancer diagnosis, ethnicity, BMI (imputed where necessary), and comorbid conditions known to influence cancer progression or thrombotic risk, including type 2 diabetes mellitus, hypertension, hyperlipidemia, hepatitis, and chronic liver disease. Cancer-related treatments, including surgery, chemotherapy, and radiation therapy, were also included as covariates. All covariates were assessed prior to the occurrence of liver metastasis. The association between heparin exposure and time to liver metastasis was evaluated using Cox proportional hazards for regression models or logistic regression models. Survival time was defined as the interval from breast cancer diagnosis to the development of liver metastasis or censoring. Multivariable Cox models were adjusted for age at diagnosis, BMI, cancer treatments (surgery, chemotherapy, radiation), and relevant comorbidities, including diabetes, hypertension, hyperlipidemia, hepatitis, and chronic liver disease. Hazard ratios (HRs) with 95% confidence intervals (CIs) were estimated. Statistical significance was defined as a two-sided p-value <0.05.

### Statistics

All statistical analyses were performed using appropriate tests as indicated for each experiment. When applicable, data were analyzed by one-way ANOVA followed by Tukey’s post hoc test, or by two-way ANOVA with Bonferroni post-tests. For non-parametric data, Kruskal–Wallis tests were applied with Dunn’s post hoc comparisons. Pairwise comparisons were conducted using Student’s t-test or Student’s t-test with Welch’s correction when variance was unequal. Correlation analyses were conducted using both Pearson and Spearman correlation coefficients. Univariate and multivariate logistic and cox regression were used to assess the occurrence of liver metastasis. All methods were applied and specified in the corresponding figure’s legends when relevant.

## Results

### E2F5 Conditional knockout tumors transplanted into the abdominal mammary gland exhibit liver tropism with patterns of invasion observed in humans

Conditional deletion of the E2F5 transcription factor in mammary epithelium (E2F5^flox/flox^ MMTV-Cre, E2F5 CKO) was previously shown to generate highly metastatic tumors, with metastases observed in the lung, lymph node, and liver ^47^. Similar to prior work, organotrophic metastasis was enriched using a serial transplantation method, but with a key difference in that cell lines were not generated at each step, nor were tumor fragments cultured *in vitro* ^8,48^. Instead, fragments of the metastatic tumors were sequentially implanted into the mammary fat pad of MMTV-Cre recipients (Figure 1A), establishing enriched lymphatic (Blue) or liver (Yellow) metastases with penetrance greater than 70% (Figure 1B).

**Figure 1.**
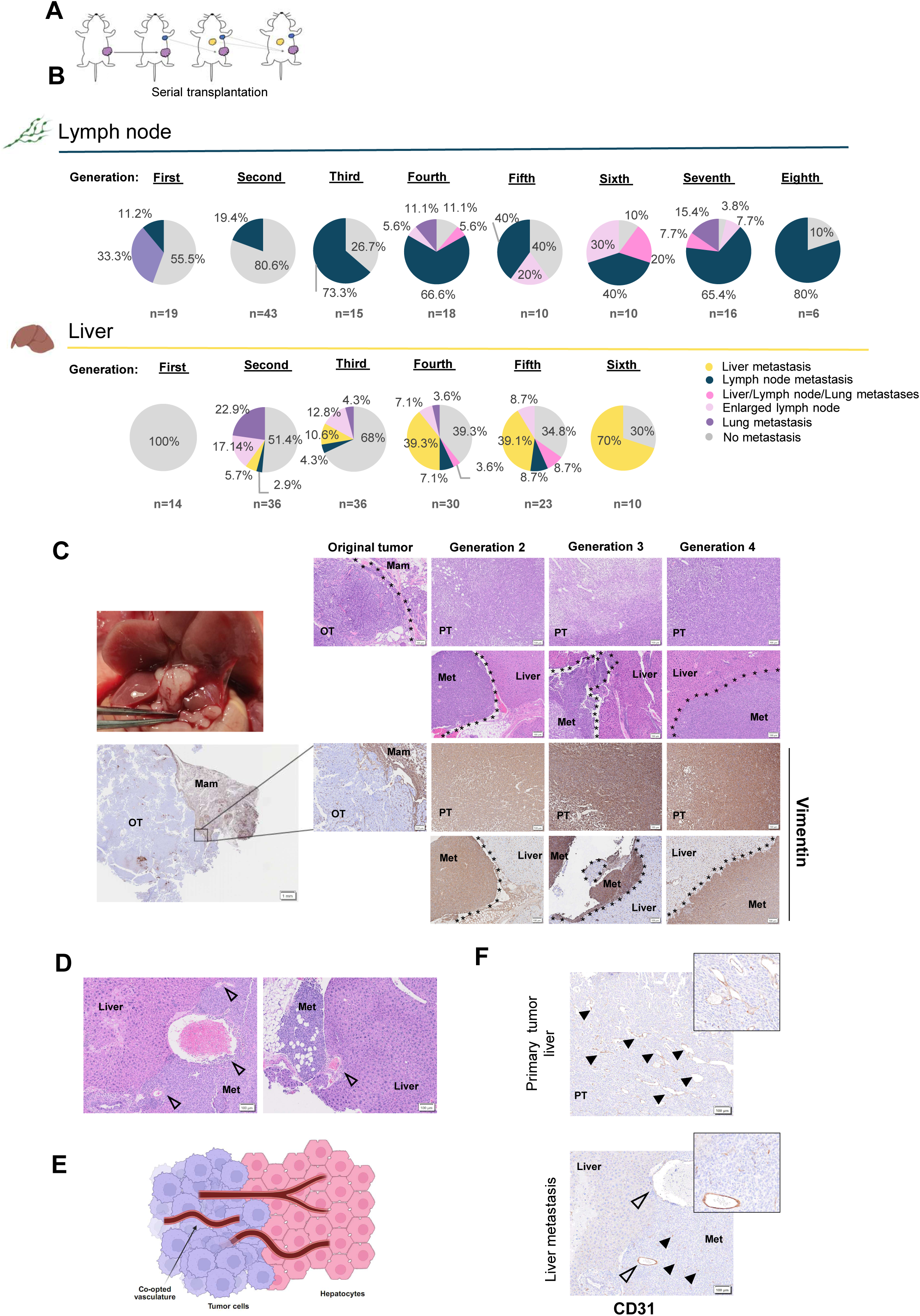
Organ tropism for liver and lymph node enhanced in E2F5 conditional knockout mammary tumors. (A) Schematic of tumor enrichment strategy. E2F5 conditional knockout mammary tumors were orthotopically transplanted into the abdominal mammary gland. Distant metastases were transplanted into the mammary glands of new recipients. Orthotopic enrichment was repeated until organ-specific metastasis was greater than 70%. (B) Organ-specific metastatic enrichment after serial tumor transplantation in lymph node (80%) and liver (70%) lineages. (C) Histological characterization of paired primary tumors and liver metastases over generations. Perivascular growth adjacent to liver lobes and subsequent parenchymal invasion in later generations was observed in liver metastasis (H&E, upper panels). Vimentin IHC on enriched primary and metastatic tumors in comparison with primary tumors (Original tumors) (Bottom panels). OT=Original tumor, PT= Enriched primary tumor, Met= Enriched metastatic tumor, LN= Lymph node. Scale bars: 100 µm and 1 mm. See also Figures S1A and S1B. (D) Replacement pattern of invasion with vessel co-option in E2F5 CKO liver metastasis. High and low caliber co-opted vessels (open arrowheads) are shown surrounded by cancer cells. Scale bars: 100 µm. (E) Schematic of the replacement growth pattern of liver metastasis characterized by the elimination of hepatocytes and vessel co-option. (F) Representative CD31 IHC showing intratumoral microvascular density (solid arrowheads) in liver metastasis compared to primary tumors. Co-opted vessels on the liver-tumor interface are indicated with open arrowheads. CD31= Platelet endothelial cell adhesion molecule-1. Scale bars: 100 µm.

Initial metastatic lesions in the liver exhibited perivascular growth, progressing to parenchymal invasion in later generations of transplantation (Figure 1C). By the second generation, liver metastases displayed increased aggressiveness and more defined EMT phenotypes characterized by elevated expression of vimentin (Figure 1C). Additional epithelial and proliferative markers were evaluated (Supplemental Figure 1A, B). Interestingly, the histology of the mouse model mimicked the human RGP for liver metastases (Figure 1D, E) based on the international consensus guidelines for scoring phenotypes of liver metastasis^26^. CD31 IHC revealed that tumors used pre-existed vasculature at the tumor-liver interface (Open arrowheads, Figure 1D, F) and presented reduced intratumoral mature vasculature (Solid arrowheads, Figure 1F).

### Transcriptome analysis uncovers enrichment of the complement and coagulation cascade in liver metastasis

To uncover organotropic drivers of liver metastasis, bulk RNA sequencing was performed on primary tumors referred to as: original primary tumors, liver and lymph node enriched primary tumors, liver and lymph node metastases and cell lines derived from these samples. Principal component analysis revealed defined clusters for the liver and lymph node tropic lineages that diverged from the primary tumors. Enriched (light red) and original liver metastases (yellow) were most closely related to normal liver samples (dark pink) (Figure 2A). Exploring the gene expression profile of liver metastases grouped by transplant generation in comparison to normal liver tissue, showed that generations 2-3 of liver metastases expressed liver-specific genes that decreased in the later generations as the gene profile became closer to the enriched primary tumors (Supplemental Figure 1C). To ensure liver metastasis samples were not influenced by potential hepatic cell contamination, a hepatocyte module core was generated (Supplemental Table 1) with genes characterizing hepatic identity and examined the core gene expression across tumor samples. Results confirmed that tumors, including liver metastases, did not approach correlation values observed in liver tissue, indicating sample purity (Supplemental Figure 1D). The top genes from the module in liver metastasis samples included hepatocyte-derived plasma proteins involved in lipid transport, coagulation and extracellular matrix regulation, suggesting an association with liver microenvironment transcriptional programs rather than structural hepatocyte markers (Supplemental Figure 1E). ssGSEA for cell type revealed a significant decrease in the hepatocyte markers signature in all tumor samples (Supplemental Figure 1F). Together, these results indicate that liver tropic tumors present a tumor-like transcriptome with modest hepatocyte-related signals.

**Figure 2.**
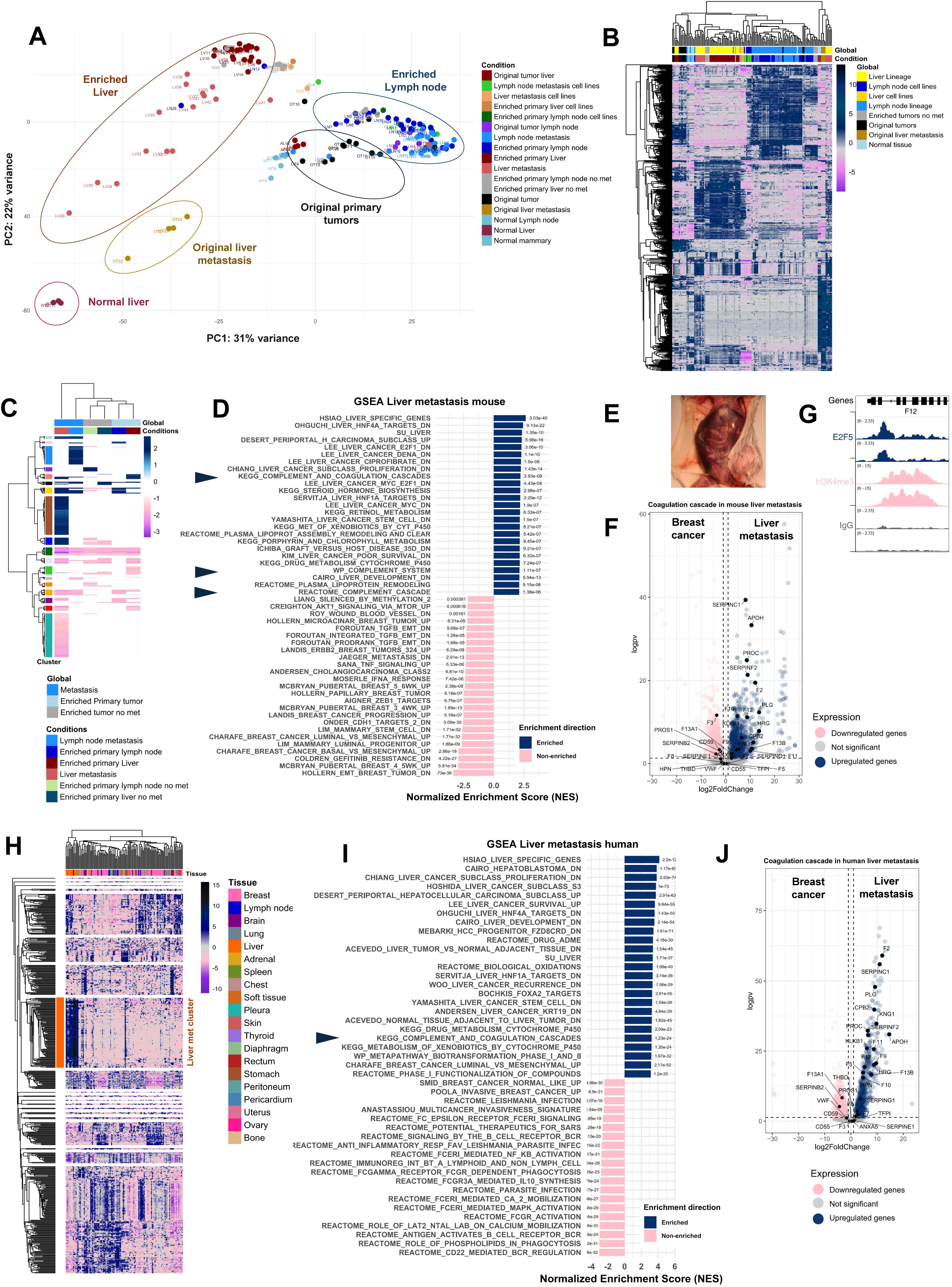
Transcriptome analysis of E2F5 CKO tumors exhibit translational relevance in human breast cancer liver metastasis. (A) Principal component analysis showing separation of clusters for Liver and Lymph Node tropic lineages. See also Figures S1C-S1F, and Supplemental Table 1. (B) Unsupervised clustering presenting the gene expression prolife of E2F5 CKO tumors and derived cell lines. (C) GSEA following enriched pathway filtering displaying unique pathway clusters per groups. See also Supplemental Table 2, and Figures S2A, S2B. (D) Top GSEA revealing the enrichment of the complement and coagulation cascade (solid arrowheads) in E2F5 CKO liver metastasis. (E) Liver metastasis bearing females developed ascites and abdominal bleeding during metastatic progression. (F) Volcano plot showing top differentially regulated coagulation genes in E2F5 CKO liver metastasis versus primary breast tumors. See also Figure S3A. (G) Integrative Genomics Viewer (IGV) showing CUT&RUN results for E2F5 and H3K4me3 occupancy at a weak promoter region of coagulation factor (F12). See also Figure S4B. (H) Unsupervised clustering on human data from the Aurora US Network (Garcia-Recio et al. 2022). See also Figure S5A. (I) GSEA highlighting the enrichment of the complement and coagulation cascade (solid arrowheads) in human breast cancer liver metastasis. See also Figures S5B-S5D. (J) Volcano plot of differentially regulated coagulation genes in human liver metastasis versus breast cancer pair tumors. See also Figures S6A and S6B.

Unsupervised clustering of the full dataset illustrated the gene expression differences between liver and lymph node lineages (Figure 2B). Given the expression of numerous genes with shared patterns, GSEA followed by filtering strategies was used to uncover mechanisms for each tropic lineage. Liver- and lymph node-enriched metastatic groups were first compared with original primary tumors, and the resulting pathways were selected only if also found when comparing metastatic tumors with enriched primary tumors. This resulted in 33 lymph- and 114 liver-metastatic pathways (Supplemental Figure 2A, B). Further comparison of all groups with original tumors defined lineage-specific clusters (Figure 2C). In total, 64 pathways were exclusively enriched in liver metastasis (Supplemental Table 2).

The top C2 curated gene sets identified by GSEA in liver metastasis (Figure 2D) revealed several expected findings, including gene sets for metastasis and breast cancer progression. However, complement and coagulation pathways were strongly enriched in the liver metastasis samples (solid arrowheads Figure 2D). This was consistent with frequent abdominal distention and bloody ascites in mice with advanced liver metastasis (Figure 2E). Differential gene expression confirmed the upregulation of coagulation genes in liver metastatic samples relative to other groups (Supplemental Figure 3A, Supplemental Table 3), including between liver metastases and original tumors (Figures 2F). F2 (Thrombin), SERPINC1, SERPINF2 and PROC showed strong upregulation in liver metastasis. Notably, among the two coagulation initiators F12 (Hageman factor), but not Tissue Factor (F3), was upregulated (Figure 2F, Supplemental Figure 3B, C, and Supplemental Table 3). F12 expression correlation analysis for liver and lymph node lineages demonstrated that expression of coagulation factors positively correlated with the expression of F12 (Supplemental Figure 3D, 4A), primarily in liver metastatic tumors.

To determine how deletion of the transcription factor E2F5 was linked to altered regulation of clotting factors in liver metastasis, CUT&RUN was performed for E2F5 in MCF-7 cells. MCF-7 was used as an independent human breast cancer model that retains endogenous E2F5 expression and presents procoagulant activity in plasma-based thrombin generation assays ^49,50^. There were defined peaks for E2F5 and H3K4m3 occupancy at a regulatory region classified as a weak promoter of F12 ^51–54^, suggesting that E2F5 regulates F12 expression (Figure 2G, Supplemental Figure 4B).

Given that our findings were in a mouse model, the significance of the top E2F5 CKO-enriched pathways in human breast cancer liver metastasis was evaluated using the AURORA US Metastasis Project data ^41^. Unsupervised clustering identified a distinct liver metastatic cluster enriched for genes regulating coagulation and lipid metabolism (Figure 2H, Supplemental Figure 5A). GSEA comparing liver metastases and primary breast tumors confirmed that the complement and coagulation cascade was enriched in human breast cancer liver metastasis (Solid arrowhead, Figure 2I) with upregulation of coagulation factors associated with the activation of the intrinsic pathway (Figure 2J) among others (Supplemental Figure 5B-D). Indeed, the key genes driving the GSEA results in our mouse model were also upregulated in human liver metastasis (Supplemental Figure 6A, B). Additionally, PAM50 molecular subtype classification of E2F5 CKO and human tumors revealed that liver metastases shared a luminal B phenotype (Supplemental Figure 6C, D).

Taken together, these results suggest that murine and human breast cancer liver metastases share expression phenotypes, with coagulation activation as part of common genetic reprogramming events.

### Tumor-derived FXII expression in E2F5 CKO liver metastatic cells was sufficient to activate coagulation *in vitro*

Coagulation factors, including FXII, were upregulated in liver metastatic lineages. To determine whether this phenotype could drive clot formation, cell lines derived from enriched primary and metastatic tumors were used to perform an *in vitro* cell-based fibrin(ogen) clotting assay (Figure 3A). This resulted in formation of fibrin(ogen) clots in the liver metastatic line (Figure 3B). To validate these structures as clots were clots, the effect of the thrombin inhibitor Dabigatran (DAB) on fibri(ogen) clot formation was evaluated ^55,56^. 24 h of DAB treatment resulted in elimination of the fibrin(ogen) network-like structure (Figure 3C). Examination of the time fibrin(ogen) clots take to form was completed in a time series. This revealed that metastatic cells from both lineages (Lymph Node and Liver) had initial clot formation after 4 h of incubation, with similar fibrin(ogen) densities. However, after 24 h, the lymphatic lineage had a more porous fibrin(ogen) network with partial fiber loss, whereas the liver metastasis cells retained more dense and less porous fibrin(ogen) networks, suggesting greater resistance to fibrinolysis (Supplemental Figure 7A).

**Figure 3.**
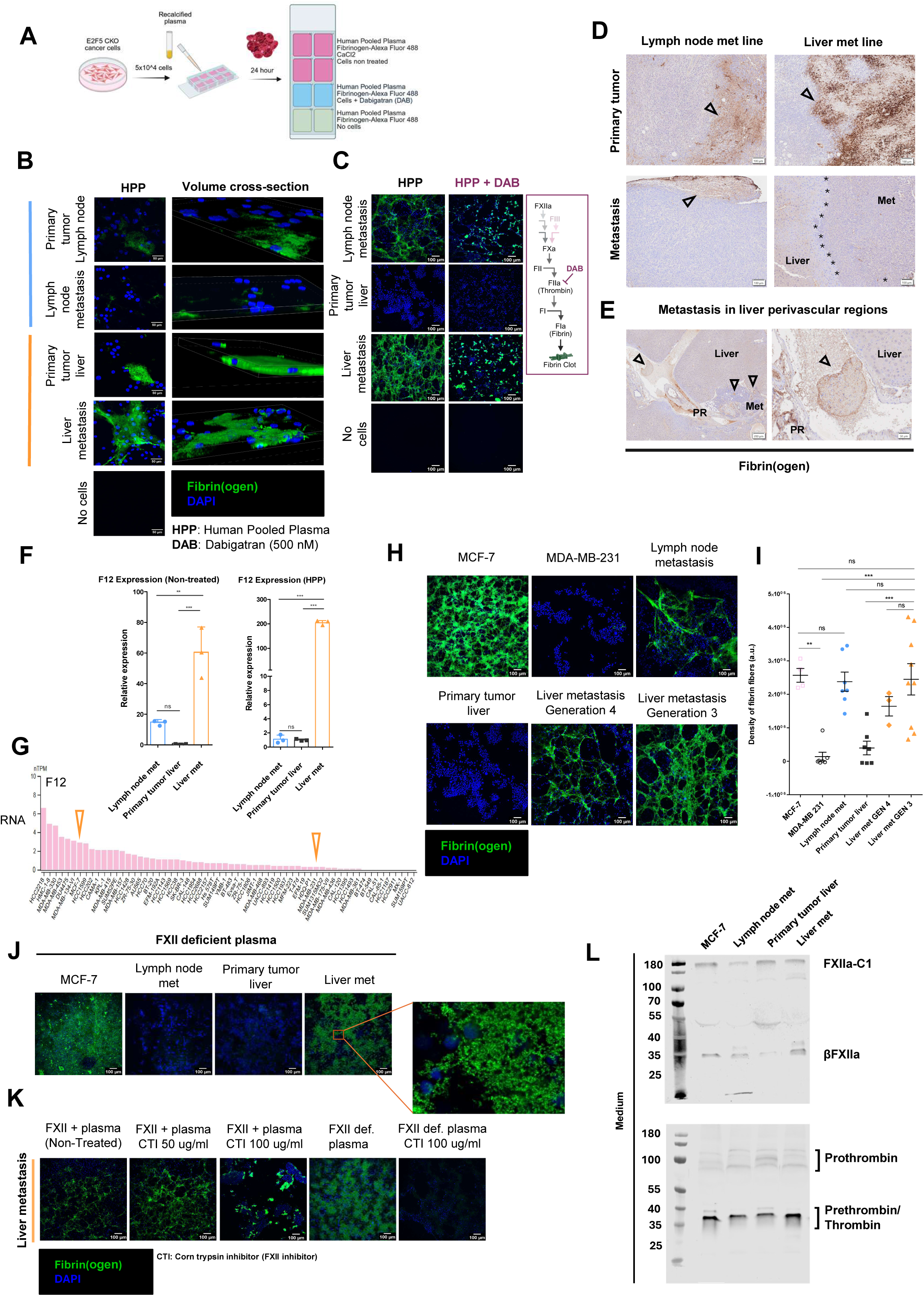
Coagulation activation is driven by tumor-derived FXII in liver metastasis. (A) Schematic of the in vitro clotting formation assay. Cells were seeded in a slide chamber and incubated with human pool plasma (HPP), CaCl2, and Alexa Fluor 488 –Fibrinogen. Conditions with/without dabigatran (DAB) were tested. (B) Representative confocal images and volume cross-section of in vitro clotting assay on E2F5 CKO tumor cell lines (n=9, scale bars: 50 µm). See also Figure S7A. (C) In vitro clotting assay on cells treated with DAB 500 nM for 24 h (Left panel, n=4, scale bars: 100 µm). Schematic representation of the coagulation cascade showing the point of inhibition of DAB and the role of thrombin in fibrin formation (Right panel). (D) Fibrin IHC identifies fibrin(ogen) fibers on primary E2F5 CKO tumors. Open arrowheads: fibrin(ogen) fibers, *: tumor-liver interface, Met: metastasis. Scale bars: 100 µm. See also Figure S7B. (E) Fibrin IHC on liver metastatic lesions growing in perivascular regions (PR). Open arrowheads identify fibrin(ogen) fibers and fibrin thrombi. Scale bars: 50 and 200 µm. (F) F12 gene expression levels in E2F5 CKO primary cell lines (Left panel). HPP treatment increases F12 expression in liver metastatic line (Right panel). Results display three independent experiments performed for each treatment and analyzed by one-way ANOVA with Tukey test post hoc. (p≤0.003=**, p≤0.0006=***, error bars: SD). (G) Relative F12 transcript abundance in human breast cancer cell lines from the Human Protein Atlas. Expression is shown as normalized transcripts per million (nTPM). Orange open arrowheads identify selected high (MCF-7) and low (MDA-MB-231) F12 expressing tumor cells. (H) Representative images comparing the *in vitro* clotting ability between human and E2F5 CKO cell lines with different F12 expression levels. Scale bars: 100 µm. Also see Figure S7A. (I) Fibrin(ogen) clot density quantification on human and E2F5 CKO cell lines. Comparison of fiber density was performed by one-way ANOVA with Tukey test post hoc (ns= not significant, p≤0.004=**, p≤0.0006=***, error bars: SEM). Dots represent individual experiments (MCF-7: n=4; MDA-MB 231, lymph node metastasis, primary tumor liver: n=7; Liver metastasis generation 4 (GEN 4): n= 3; Liver metastasis generation 3 (GEN 3): n=9). (J) In vitro clotting assay performed with FXII-deficient plasma on cells with high (MCF-7 and liver metastasis) and low (Lymph node metastasis and primary tumor liver) F12 expression (n=4. Scale bars: 100 µm). Quantification of fibers’ density on multiple experiments can be found in Figure S8A. (K) Effect of FXII inhibitor CTI at 50 and 100 µg/mL on clotting formation on liver metastatic cells incubated with FXII+ plasma (n=3) or FXII deficient plasma (n=4). Scale bar: 100 µm. Quantification of fiber density in multiple experiments can be found in Figure S8B. (K) Representative immunoblots of FXIIa and Thrombin active forms in liver metastatic cell conditioned medium. Densitometry of bands normalized by total protein are represented as dots and bar plots in Supplemental Figure S8D. See also protein expression over time and non-cells control in Figure S8E.

In mouse tumor samples, fibrin(ogen) was analyzed by IHC, revealing dense fibrin(ogen) fiber depositions in necrotic areas of enriched primary tumors from the liver lineage. Unexpectedly, liver metastases had no fibers in the metastatic core or at the tumor-liver interface (Figure 3D). This was supported by the absence of the fibrinogen gamma subunit in liver metastasis, which is essential for the interaction of fibrinogen with FXIII-B and thrombin-mediated cleavage during fibrin polymerization ^57^ (Supplemental Figure 7B). In the perivascular regions of the liver, fibrin(ogen) staining was observed surrounding tumor cells associated with fibrin thrombi in blood vessels (Figure 3E), suggesting that the fibrin clot was a component of circulating tumor cells.

After observing fibrin(ogen) fibers in the tumors, the expression of F12 in primary cell lines derived from these E2F5 CKO tumors was tested. A significant upregulation of F12 in the liver metastasis lines was noted relative to lymph node metastasis and primary tumor lines. Liver metastatic cells incubated with human pooled plasma (HPP) displayed a dramatic upregulation of F12 (Figure 3F). To evaluate whether F12 expression was associated with fibrin(ogen) clot formation in human breast cancer models, F12 RNA expression was surveyed in human breast cancer cell lines. MCF-7 had ∼12 fold higher F12 transcript levels relative to MDA-MB-231; they were selected as F12-high and F12-low controls for in vitro clotting assays (Cell line - F12 - The Human Protein Atlas from v24.1.proteinatlas.org) (Figure 3G). Testing these cell lines in the *in vitro* fibrin(ogen) formation assay demonstrated that MCF-7 cells, with high expression of F12, exhibited more dense fibrin(ogen) networks, whereas MDA-MB-231 resulted in no fibers being detected (Figure 3H, I and Supplemental Figure 7A).

To test whether tumor cell derived FXII was sufficient to drive fibrin(ogen) clot formation, the clotting assay was repeated with FXII deficient plasma. Cells expressing high levels of F12 formed fibri(ogen) networks, demonstrating that it was sufficient to drive fibrin clot formation (Figure 3J, Supplemental Figure 8A). Further, the effect of corn trypsin inhibitor (CTI), a potent FXII inhibitor, was tested in the formation of fibrin(ogen) fibers by liver metastatic cells. Plasma with or without FXII and with increasing concentrations of CTI were tested. This revealed the elimination of fibrin nets upon incubation with CTI at 100 µg/mL (Figure 3K, Supplemental Figure 8B). To assess the contribution of the extrinsic clotting pathway, liver metastatic cells were treated with tissue factor pathway inhibitor (TFPI) at 0.001 mg/mL. No significant differences in fiber formation were observed after treatments (Supplemental Figure 8C). Immunoblots confirmed elevated βFXIIa and Thrombin active forms on liver metastatic cells conditioned medium (Figure 3L, Supplemental 8D, E) and βFXIIa on cell lysates (Supplemental Figure 8F), results consistent with ongoing activation of the coagulation pathway. Notably, bands corresponding to βFXIIa in complex with the esterase inhibitor C1 ^58^ were observed. FXIIa IHC revealed elevated intratumoral FXIIa expression in liver tropic tumors as well as their derived cell lines (Supplemental Figure 8G). Collectively, these results suggest that metastatic cancer cells with liver tropism exhibit a procoagulant phenotype associated with elevated expression of FXII. In this context, tumor-derived FXII was sufficient to initiate coagulation in E2F5 CKO liver metastases.

### Glutamine deprivation induces mechanisms of FXII activation and coagulation

Plasma FXII zymogen is activated through the interactions with negatively charged surfaces ^59^. Externalization of the anionic phosphatidylserine (PS) occurs in cancer cells due to metabolic stress ^60^, and has been associated with pro-coagulant phenotypes in triple-negative breast cancer ^61^. Thus, similar metabolic and membrane alterations could induce the activation of FXII and formation of clots in E2F5 CKO tumor cells. E2F5 CUT&RUN on MCF-7 and HC11 cells revealed E2F5 target genes involved in lipid and glutamine metabolism with direct impacts on redox homeostasis, including glutaminase (GLS), acyl-CoA Oxidase 1 (ACOX1), and Fatty acid synthase (FASN) (Figure 4A).

**Figure 4.**
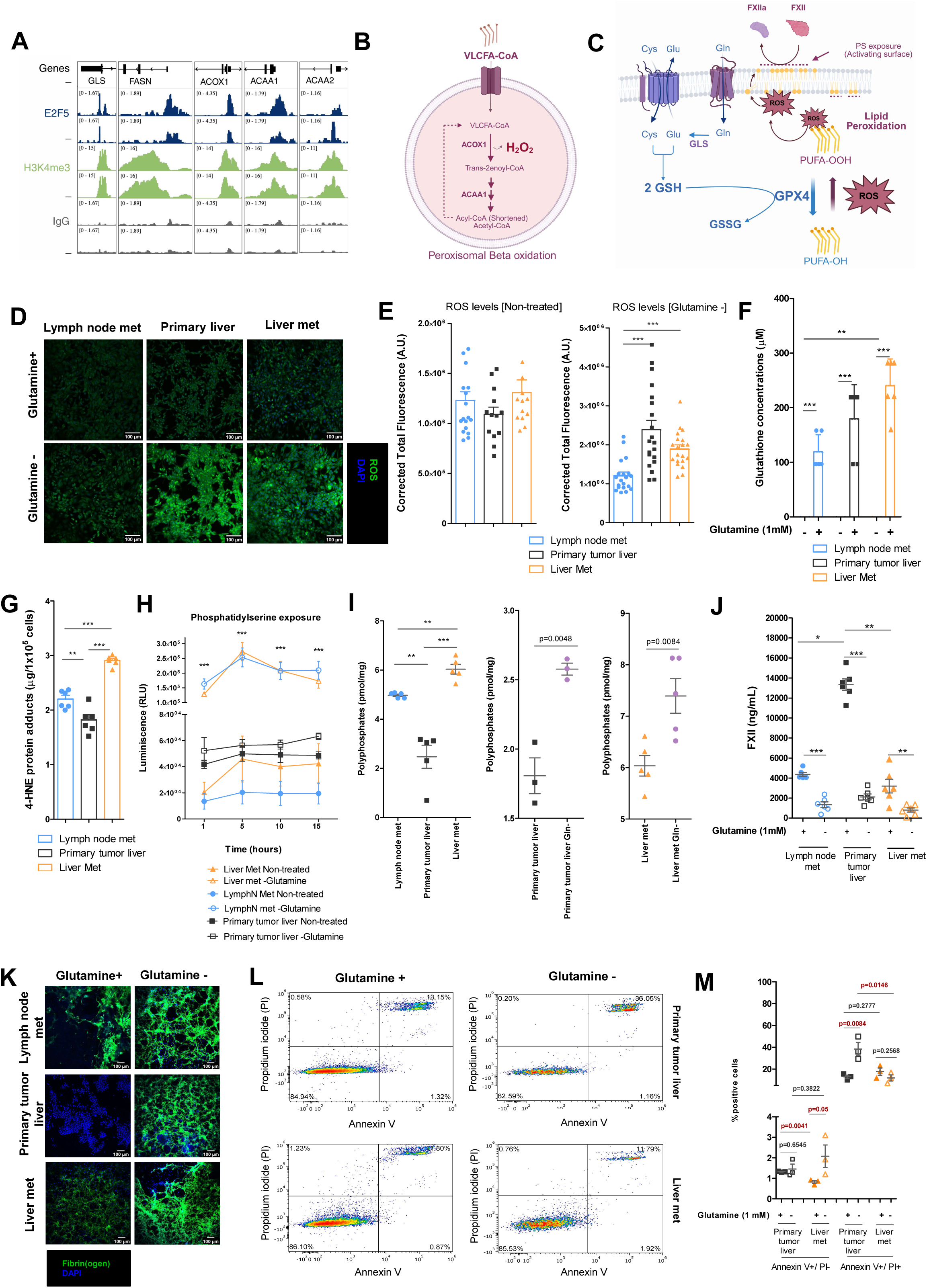
Cell stress and lipid peroxidation levels control FXII activation through E2F5 regulated mechanisms. (A) Integrative Genomics Viewer (IGV) showing CUT&RUN with HC11 cells for E2F5 and H3K4me3 occupancy at the promoter of genes Glutaminase (GLS), Fatty acid synthase (FASN), Peroxisomal acyl-coenzyme A oxidase 1 (Acox1), Acetyl-CoA acyltransferase 1 (ACAA1), and 2 (ACAA2). (B) Illustration of H_2_O_2_ production during peroxisomal β oxidation. (C) Model of FXII activation driven by phosphatidylserine externalization due to elevated lipid peroxidation activity in liver metastatic cells. (D) Representative images of overall ROS signal using confocal microscopy in primary and metastatic E2F5 CKO cell lines growing in glutamine-replete and glutamine-deprived media. Scale bars: 100 µm (E) ROS signal quantification. Dots represent biological replicates. Statistical analysis was performed by Kruskal-Wallis with Dunn post hoc, p≤0.001=***, error bars: SEM. (F) Glutathione (GSH) quantification in primary and metastatic E2F5 CKO cell lines growing in glutamine-replete and glutamine-deprived media. Statistical analysis was assessed by one-way ANOVA with Tukey test post hoc (n=5; p≤1.8×10-3=**, p≤1.3×10-4=***; error bars: SD). (G) Quantification of 4-hydroxynonenal (4-HNE)-protein adducts, a lipid peroxidation waste product, in E2F5 CKO cell lysates using competitive ELISA. Differences in concentration were analyzed by one-way ANOVA with Tukey post hoc (n=6, p≤4.5×10-3=**, p≤1.1×10-5=***, error bars: SEM). (H) Annexin V kinetics on E2F5 CKO cell lines to identify externalized phosphatidylserine (PS) signal in E2F5 CKO cells growing in glutamine-replete and glutamine-deprived media (H). Results are representative of three individual assays analyzed by two-way ANOVA with Bonferroni post-tests (p<0.001= ***, error bars: SEM). (I) Concentrations of FXII activator polyphosphate (PolyP) were determined on E2F5 CKO cell lines (Left panel). Samples were pre-treated with proteinase K, RNase, and DNase, and agarose electrophoresis was performed to check for any residual RNA and DNA. Total PolyP was quantified as AxD/(Wx(Vt/Va)), where A: Represents interpolated amounts of PolyP from standard curve in pmol, D: The sample dilution factor, W: the cell pellet weight in mg, Vt: the total sample volume, and Va: the volume of sample loaded into the well. Results are expressed in pmol/mg and Kruskal-Wallis with Dunn post hoc was used for statistical analysis (n=5; p≤0.01=**, p≤0.001=***, error bars: Mean and SEM). PolyP concentration following Glutamine-depravation (right panels). Differences were calculated with Student T test, error bars: Mean with SEM. (J) Concentration of FXII zymogen in E2F5 CKO cell conditional media determined by ELISA. Results were reported in ng/mL and analyzed by Student T test (n=6, p≤0.01=*, p≤0.007=**, p≤0.0001=***, error bars: Mean with SEM). (K) Representative in vitro clotting assay with E2F5 CKO cells cultured in glutamine-replete and glutamine-deprived media for 24 h (n=3, scale bars: 100 µm). (L) Representative flow cytometry results on primary tumor cells and liver metastatic cells cultured in glutamine-replete and glutamine-deprived media for 24 h. Percentage of early (Annexin V+/Propidium iodide (PI-)) and late (Annexin V+/Propidium iodide (PI+)) apoptotic cells were measured using FlowJo. Assay was performed twice with three biological replicates per condition. (M) Flow cytometry statistical analysis was performed by Welch’s t test, error bars: Mean with SEM.

ACOX1-driven β oxidation generates hydrogen peroxide (H_2_O_2_) as a by-product ^62^, and its dysregulation increases ROS levels ^63^ (Figure 4B). In parallel, GLS is essential to de novo GSH synthesis ^24^, required to control ROS ^64^. Disruption of this balance leads to ROS accumulation, promoting polyunsaturated fatty acid (PUFA) peroxidation, membrane damage ^65^ and phosphatidylserine (PS) externalization ^66^. Exposed PS may serve as an activating surface for FXII zymogen (Figure 4C). Therefore, we hypothesized that E2F5 CKO liver metastatic cells present a procoagulant phenotype due to alterations in glutamine and lipid metabolism associated with E2F5 loss.

To test whether these cells showed altered redox responses to glutamine availability, cells were cultured in glutamine-deprived medium, which revealed elevated ROS in enriched primary and liver metastatic lines relative to the lymph node metastatic line (Figure 4D, E). Consistent with these observations, GSH concentrations were measured in media with or without glutamine. Liver-tropic lines showed significantly higher GSH under replete conditions (Figure 4F). Next, evidence of lipid peroxidation was tested under glutamine-replete conditions by measuring the concentrations of 4-hydroxynonenal (4-HNE) protein adducts ^67^. Liver metastatic cells showed elevated 4-HNE adducts levels (Figure 4G), indicating increased lipid peroxidation under basal conditions. Together, these findings suggest that liver metastasis sustains elevated lipid peroxidation while maintaining GSH buffering capacity, consistent with a redox-adapted state that may allow these cells to tolerate oxidative stress. After establishing elevated lipid peroxidation in liver metastasis, lipid availability and storage were examined to determine whether lipid alterations could induce the oxidative phenotype observed. Intracellular free fatty acids (FFA) were quantified, revealing elevated FFA levels in metastatic cells compared to primary tumor liver selected for liver tropism (Supplemental Figure 9A). Interestingly, IHC for Perilipin 2 (PLIN2), a constitutive marker of lipid droplet (LD) density ^68^, revealed that liver metastatic tumors lacked detectable LDs in most tumor regions (Supplemental Figure 9B). However, this contrasted with the elevated expression of the mature membranous variant of the LDL receptor (LDLR) ^69^ in a liver metastatic cell line (Supplemental Figure 9C). These cells also showed elevated FASN, a key enzyme in *de novo* FA synthesis in cancer ^70^ (Supplemental Figure 9C), compared to lymph node-metastatic cells. These results suggest that liver metastatic cells are susceptible to lipid peroxidation due to increased lipid uptake, enhanced de novo lipid synthesis, and defective LD formation. Impaired LD formation might reduce the antioxidant capacity of liver-tropic cells ^71–73^.

The implications of elevated oxidative stress for FXII activation were investigated (Figure 4C). First, PS externalization on the plasma membrane was assessed in E2F5 CKO cells using an Annexin V assay. Glutamine removal on exposed PS levels was evaluated, revealing a sharp rise of externalized PS on metastatic cell lines upon glutamine deprivation without significant changes in enriched primary tumor cells (Figure 4H). Additionally, liver metastatic cells exhibited higher concentrations of Polyphosphates (PolyP), an activator of FXII, compared with enriched primary tumor and lymph node metastasis lines (Figure 4I). Glutamine deficient medium also increased the concentrations of PolyP in liver-tropic cells (Figure 4I).

Based on these findings, we hypothesized that FXII activation is enriched in glutamine deficient conditions. To test this, an ELISA for FXII zymogen, the inactive form of FXII, was perfomed in conditioned media. As expected, the concentration of FXII zymogen decreased in glutamine-deprived medium for all cell lines. Notably, primary tumor cells exhibited elevated FXII zymogen concentrations under glutamine-replete conditions (Figure 4J), suggesting fewer basal FXII activation events compared with pro-coagulant metastatic cells. Upon glutamine deprivation, each of the lines exhibited dense fibers in the *in vitro* clotting assay (Figure 4K). Therefore, the effect of glutamine deprivation on PS exposure was further examined by flow cytometry. Under glutamine deprivation, primary tumor lines showed an increment of late apoptotic cells (Annexin V+/Propidium iodide (PI+)) compared to liver metastatic cells. The percentage of late apoptotic cells in liver metastatic cells remained unchanged between conditions (Figure 4L, M). These observations indicate that the levels of cell stress required for fibers formation on the primary tumor line exceeds the threshold compatible with cell survival, as cells reach late apoptotic states characterized by irreversible death.

Together, these findings suggest that altered redox and lipid peroxide metabolism in liver metastatic cells is associated with increased PS externalization and PolyP availability, creating conditions that may favor FXII activation and fibrin(ogen) clot formation.

### Immune microenvironment characterization in pro-coagulant tumors

In examining the RNAseq data, together with an upregulation of coagulation/fibrinolysis factors, liver metastatic tumors exhibited overexpression of complement, acute-phase inflammatory response, and neutrophil granule serine-protease genes (Figure 5A), characteristic of inflammatory response in infection diseases and cancer ^74,75^. Gene set variation analysis (GSVA) suggested an innate immune activation on liver metastatic tumors with enhancement of neutrophil granule, macrophage and IL-1β pro-inflammatory signatures (Figure 5B). Furthermore, a ssGSEA analysis for immune cell type was performed, and a distinctive macrophage signature in enriched liver tropic tumors was noted (Figure 5C and Supplemental Figure 10A). F4/80 IHC confirmed the elevated macrophage infiltration in these groups (Supplemental Figure 10B), and notably, the inflammation-related cyclooxygenase (COX)-high macrophage signature was also elevated (Figure 5D). It is interesting to note that FXIIa was reported to induce macrophage secretion of TNF-α, IL-6, IL-12, and IL-1β in mice ^39^. Aligned with these observations, TNF-α expression was elevated in liver tropic tumors (Supplemental Figure 10C). Moreover, the M2 macrophage polarization signature was elevated in liver tropic tumors, and this positively correlated with elevated coagulation signature activity (Figure 5E). The impacts on fibrin(ogen) clot formation with the macrophage-secreted cytokines TNFα and IL-1β upon FXII induction was tested. Previous studies reported TNF-mediated FIII upregulation ^76^ and the alteration of blood clots upon IL-1β action ^77,78^. Here, results demonstrated modulation in the clotting activity of cancer cells incubated with both cytokines (Supplemental Figure 10D). In the context of FXII capacity to stimulate macrophages derived production of these pro-inflammatory cytokines, these data support the plausibility of FXII-macrophage-cytokine-coagulation crosstalk on liver tropic tumors.

**Figure 5.**
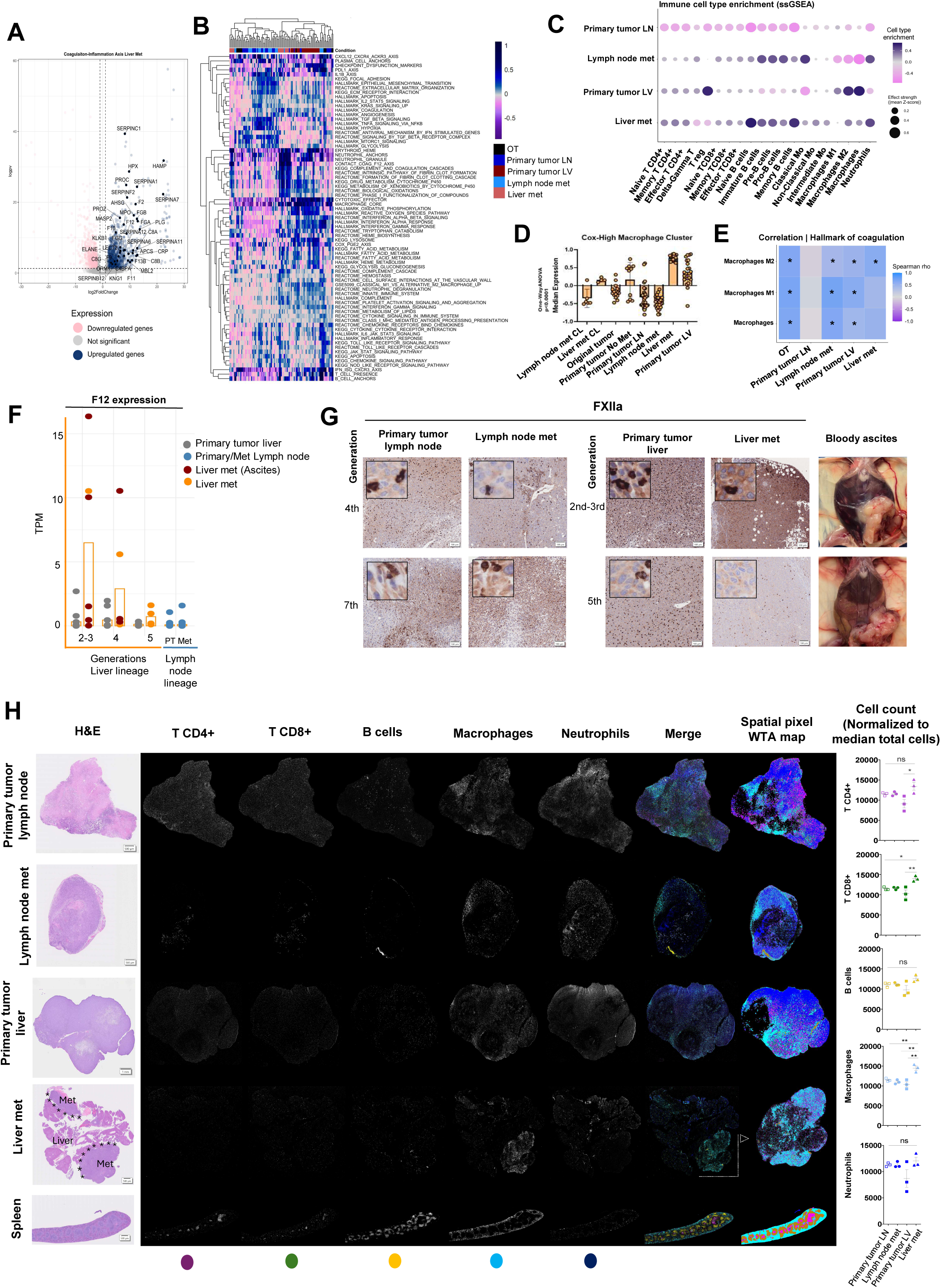
Characterization of tumor immune microenvironment and its association with coagulation. (A) Volcano plot from DESeq showing the upregulation of complement and inflammatory response in liver metastatic tumors. (B) GSVA exploring immune signatures in E2F5 CKO tumors. (C) Identification of immune cell type enrichment by ssGSEA in E2F5 CKO tumors. Cell markers were selected based on PanglaoDB - A Single Cell Sequencing Resource For Gene Expression Data and ScTypeDB -sc-type. Also see Figure S10A, S10B. (D) Cox-High macrophage signature median expression across samples. Statistical analysis was performed using one-way ANOVA (p≤0.0001, error bars: SD). (E) Correlation between macrophage signatures and coagulation activity in E2F5 CKO tumors (Spearman correlation, p≤0.05=*). (F) Relative F12 transcript abundance in liver tropic tumors across generations of transplantation. Expression is shown in transcripts per million (TPM). Liver metastatic tumors from females with bloody ascites were highlighted in red. Lymph node tropic tumors (blue) were used as controls. (G) FXIIa IHC showing the expression of intratumoral FXIIa across transplant generations for both lymph node and liver lineages (Left panel, scale bars: 100 µm). Representative image showing the incidence of abdominal bleeding and ascites on liver metastasis bearing females on generation two versus five (Right panel). Also see Figure S10E. (H) Representative image mass cytometry (IMC) and immune cell quantification in primary and metastatic E2F5 CKO tumors. H&E are shown in left panel (*: Tumor-liver interface, scale bars: 500 µm and 1mm). Raw IMC signal per marker with merge-colored images, and spatial pixel winner takes all (MTA) maps generated with Iolite output matrices are shown in the center panel (Scale bars: 500 µm and 1mm). Immune cells count normalized to median total cells, and statistical comparison across samples are shown in the right panel. Statistical analysis was performed by one-way ANOVA with Tukey test post hoc (n=3; error bars: Mean with SEM; T CD4+ (Magenta): p≤0.015=*, T CD8+ (Green): p≤0.042=*, p≤0.0047=**, B cells (Yellow), Macrophages (Cyan): p≤0.0014=**, and Neutrophils (Blue)). Also see Figure S11 and S12.

F12 expression exhibited gradual reduction over generations of transplantation (Figure 5F) and correlated with reduced protein levels in liver tropic tumors (Left panel, Figure 5G). These observations also correlated with a marked depletion of ascites in liver metastasis bearing females at the latest generation of transplantation (Right panel, Figure 5G). Moreover a reduction of macrophages infiltration in tumors with low FXII expression was noted (Supplemental Figure 10E). Based on this, the E2F5 CKO tumor immune landscape was explored with imaging mass cytometry (IMC) analysis and the populations of T lymphocytes (CD4+, CD8+), B lymphocytes, neutrophils, and macrophages were quantified with IMC, and flow cytometry (Supplemental Figure 9F). IMC results were processed following the MeXpose pipeline ^79^ with some modifications. The intensity per marker/per area was determined for each tumor (Supplemental Figure 11). These values were assigned after cell segmentation to classify cell types for quantification and image construction (Supplemental Figure 12). Results revealed elevated macrophage infiltration and considerable T CD4+ and CD8+ cell recruitment in liver metastatic tumors (Figure 5H). Notably, flow cytometry analysis demonstrated a reduction of T CD8+ population expressing the co-stimulatory molecule CD28 in liver metastasis (Supplemental Figure 10F).

Together, these results suggest a role of tumor-derived FXII in the generation of a pro-thrombotic state in liver metastatic bearing females that favor the recruitment of macrophages during the initial stages of liver colonization.

### Low molecular weight heparin treatment reduces liver tropism and metastatic burden

In the serial transplantation (Figure 1), liver tropic lines were implanted into the abdominal mammary gland and metastases were routinely observed in a perivascular location adjacent to the intrahepatic vena cava and portal vein. Blood flow from the abdominal mammary gland involves both the thoracoepigastric and caudal epigastric veins. These capillaries connect with the inferior vena cava (IVC) and femoral vein respectively ^80,81^. Both routes pass through the intrahepatic region of the IVC before reaching the lungs for oxygenation (Figure 6A). Based on vascular flow, we hypothesized that the location of the transplanted tumors impacts organotropism. Liver tropic E2F5 CKO tumors were implanted into the left thoracic mammary glands (#2/3) and compared the metastatic progression with the same tumors implanted into the abdominal (#4) mammary gland. The thoracic implantation resulted in a 92% reduction of liver tropism and 111% increase in tumor-bearing mice that lacked metastases relative to abdominal implantations (Figure 6B, C). Circulating tumor cells (CTCs) colonies were identified by collecting blood through cardiac puncture and then culturing the CTCs, observing a slight increase of CTC colonies in mammary gland 4 transplanted females (Supplemental Figure 13A). These results indicate that liver metastatic cells have become specialized for liver colonization. Altering the vascular route may expose liver-tropic CTCs to the small capillary beds in the lungs, resulting in pulmonary filtration in the lung immune microenvironment and may have ultimately limited seeding in the liver.

**Figure 6.**
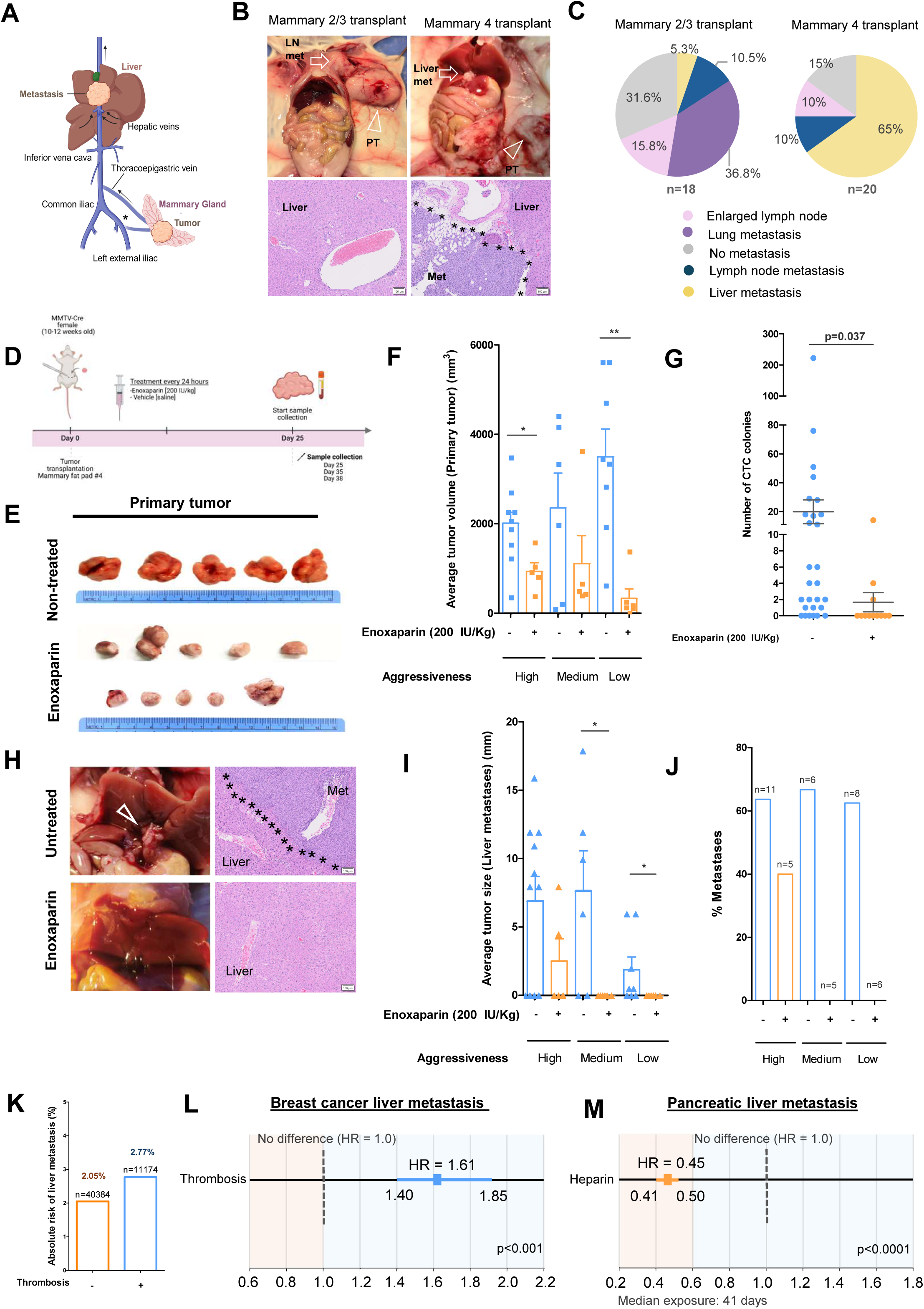
Low molecular weight heparin treatment and vascular route alterations reduce liver tropism and metastatic burden. (A) Illustration of systemic venous circulation from the point of tumor implants on abdominal mammary gland to the lungs passing through the common iliac across the center of the liver and highlighting the recurrent position of metastatic lesions observed on E2F5 CKO model (*: caudal epigastric vein). (B) Representative images of liver metastatic tumor implantation into the thoracic (2/3) or abdominal (4) mammary glands (open arrowheads) with the incidence of metastases (Upper panel, white arrows), and H&E showing metastatic tumors invading liver lobes (Bottom panels, *: tumor-liver interface, scale bars: 100 µm). (C) Percentage of metastases observed on transplanted females in Panel B. (D) Schematic representation of enoxaparin *in vivo* treatments. Liver metastatic tumors (Low, medium and high aggressive) were implanted on the abdominal mammary gland of 10- to 12-week old MMTV-Cre females and treatments with enoxaparin 200 IU/Kg or vehicle (Saline) were applied every 24 h until control group’s primary tumors reached 2000 mm. Blood, and tumors were collected for further analysis. (The low, medium, and high aggressive categories were based on the pathological signs of hemorrhagic ascites observed in liver metastasis-bearing females). (E) Representative images of primary tumors from high and low aggressive groups (Scale in cm). (F) Average primary tumor volume (mm^3^). Each tumor is represented by a dot in a bar plot. Kruskal-Wallis with Dunn post hoc was used for statistical analysis (p≤0.01=**, p≤0.05=*, error bars: Mean with SEM). (G) Circulating tumor cells (CTC) colonies were identified by cardiac puncture followed by blood fractionation and colony formation assay experiments. Each dot represents the total number of CTC colonies per mouse. Student T test with Welch’s correction was applied for statistical analysis (error bars: Mean with SEM). (H) Representative images of liver metastasis location and histology(Open arrowhead indicates liver metastatic tumors, *: tumor-liver interface, scale bars: 100 µm). (I) Average size of liver metastases per mouse (mm). Each tumor is represented by a dot in a bar plot. Statistics were assessed by one-way ANOVA with Tukey post hoc, and Kruskal-Wallis with Dunn post hoc, p≤0.01=**, p≤0.05=*, error bars: Mean with SEM). (J) Incidence of metastasis in transplanted females. (K) Retrospective cohort studies in humans exploring the absolute risk of developing liver metastasis in breast cancer patients with thrombosis. (L) Adjusted multivariable cox proportional hazards model showing the hazard of liver metastasis in breast cancer patients with thrombosis. (M) Univariate logistic regression indicating the incidence of liver metastasis on heparin-exposed pancreatic cancer patients.

Considering all the above data, liver tropic tumor cells activate components of the coagulation cascade and promote fibri(ogen) deposition. This activity may facilitate the formation of fibrin-rich structures that protect cells from adverse conditions in circulation and favor their lodging within the hepatic vasculature. To test this hypothesis, mice that had been implanted with liver tropic tumor lines were treated using low molecular weight heparin (enoxaparin) and assessed whether liver metastasis was impacted.

Three liver metastatic tumor lines classified as high-, medium-, or low-aggressive based on observable pathological signs of ascites were transplanted into the mammary fat pad. Treatment with enoxaparin 200 IU/kg was applied every 24 h until primary tumors reached endpoint in non-treated groups (Figure 6D). Heparin had a modest impact on primary tumor growth in high and medium aggressive tumors (Supplemental Figure 12B) while the low aggressive group exhibited an obvious reduction in primary tumor growth (Figure 6E, Supplemental Figure 13B) (Two-way ANOVA, p≤0.01=**, p≤0.001=***). Tumor volume measurements at endpoint indicated significant reduction in high and low aggressive groups (One-way ANOVA, Tukey post hoc and Kruskal-Wallis, Dunn post hoc, p≤0.01=**, p≤0.05=*) (Figure 6F).

To test the effect of heparin on liver metastasis, circulating tumor cells were examined by collecting blood at endpoint and performing colony formation assays. CTC colonies were significantly decreased in heparin-treated groups (Figure 6G). Liver metastases were also examined across all groups and heparin treatment eliminated metastasis for the medium and low aggressive metastatic lines. In the highly aggressive group, the incidence of metastasis decreased from 64% in control mice (n=7 of 11) to 40% in heparin-treated mice (n=2 of 5) (Figure 6H, I, J). (One-way ANOVA, Tukey post hoc and Kruskal-Wallis, Dunn post hoc, p≤0.01=**, p≤0.05=*). The metastatic lesion sizes displayed a modest positive correlation with the number of CTC colonies in non-treated groups (Supplemental Figure 13C, D).

The role of the intrinsic clotting cascade in human liver metastasis was determined by retrospective cohort studies using de-identified electronic health record (EHR) data from the Truveta platform. The association between thrombosis and time to liver metastasis using Cox proportional hazards regression models was explored. Among the 57,529 breast cancer patients included in the analysis, 11,174 (19.4%) experienced a thrombotic event during follow-up, while 40,384 had no documented thrombosis. Liver metastasis occurred in 310 patients (2.77%) in the thrombosis group and in 830 (2.05%) in the non-thrombosis group (Figure 6K). In multivariable Cox proportional hazards models adjusting for demographic factors, comorbidities, and cancer treatments, the presence of thrombosis was associated with a 61% higher hazard of liver metastasis compared with patients without thrombosis (HR 1.61; 95% CI, 1.40–1.85; p < 0.001) (Figure 6L). This association remained robust after adjustment for all covariates included in the model.

Next, the effect of anticoagulant treatment on the development of liver metastasis was investigated. Heparin-treated breast cancer patients had a median exposure duration of three days with four doses over time. The short-term drug exposure observed did not allow us to determine the impact of the drug on liver metastasis prevention. Consequently, the effect of heparin in pancreatic cancer patients was examined. Pancreatic cancer exhibits a high incidence of liver metastasis ^82–84^ and is the malignancy with the highest rate of VTE ^85,86^. The median heparin exposure of the pancreatic cancer cohort was 41 days with approximately six doses administered over time. Univariate logistic regression showed that subjects exposed to heparin had a significantly lower hazard of developing liver metastases (odds ratio: 0.52; p: < 0.0001), indicating a 48% reduction compared to those not exposed. The multivariable Logistic model-adjusted for demographic, clinical, and comorbidity variables showed a similar result, with heparin exposure associated with a 55% lower hazard of liver metastasis (HR = 0.45; 95% CI, 0.41–0.50 p: < 0.0001) (Figure 6M).

Together, these results suggest a protective role for a pro-thrombotic environment in the migration and invasion of breast cancer liver metastasis and indicate that targeting the coagulation cascade may reduce liver metastasis incidence in humans.

## Discussion

Here the development of liver-specific metastatic breast cancer tumor lines and tumor cell lines derived from E2F5 CKO mice are reported and compared with lines enriched for lymphatic metastasis. Importantly, this enrichment was completed in syngeneic immune competent mice and used tumor fragments. Primary cell lines generated from tumors maintained their liver tropism after mammary fat pad injections. This important new model revealed a critical role for the coagulation system in liver metastasis. Indeed, our data suggested that E2F5 normally represses F12 expression, as tumor cells from CKO mice exhibited elevated FXII transcript and protein levels. This expression, combined with metabolic alterations, resulted in tumor cells that had the ability to drive fibrin clots and induce macrophage recruitment. In a key finding, blocking clotting with low molecular weight heparin resulted in suppression of liver metastasis. The translational relevance of these results was supported by retrospective cohort studies using human data, which indicated that thrombosis was associated with increased incidence of liver metastasis in breast cancer patients.

Recent reports have highlighted the role of elements of the coagulation system in cancer metastasis. Aspirin showed successful anti-metastatic properties by limiting the availability of platelet-derived thromboxane A2 involved in the induction of immunosuppressive T cells ^37^. Likewise, a recent study showed that tumor cells form heteroaggregates with platelets through the generation of thrombin and fibrin polymerization ^87^. *In vitro* coagulation assays in human plasma have demonstrated that increased concentrations of FXII augmented fibrin(ogen) fiber density ^88^ and less permeable fibrin clots have been related to patients with venous thromboembolism ^89,90^. Concordantly, it was reported that FXII haploinsufficiency confers protection against VTE in humans ^91^. This extensive study summarized years of investigation ^92–96^ confirming the role of FXII in pathological thrombosis. Notably, FXII reciprocally activates the kallikrein-kinin-bradykinin system ^97^ which induces inflammation and vascular leakage, a process that may facilitate metastatic cell extravasation ^98,99^ and reinforcing a pro-tumorigenic function for FXII.

The intrinsic pathway of coagulation starts with the activation of FXII zymogen upon interaction with negatively charged surfaces ^97,100^. Different cancer-relevant molecules serve as FXII activators, including phosphatidylserine ^61,101^ and Polyphosphate ^102^. Both factors are related to oxidative stress and redox homeostasis ^66,103^. Cancer cells have elevated production of ROS and accumulate lipids ^64^. This metabolic stress can be induced by oxygen or nutrient deprivation, and lipid metabolism is particularly affected. Hypoxia induces changes in the use of carbon sources and promotes fatty acid uptake ^104^. Given that nearly three quarters of the blood that circulates in the liver is non-oxygenated venous blood, the liver is an oxygen-limited organ ^105^. This liver hypoxic environment could exacerbate the dependencies on lysosome function for the uptake of external lipids, creating a high oxidative microenvironment ^64^, metabolically affecting breast cancer cells that usually depend on external supplies to maintain lipid homeostasis ^106^.

The requirement of ROS scavenging by GSH as an essential metabolic adaptation in breast cancer liver metastasis has been reported ^24^ and glutamine contributes to the pull of glutamate necessary for the novo biosynthesis of GSH. This cofactor is essential for the reduction of oxidized polyunsaturated fatty acids (PUFA-OOH) and the control of lipid peroxidation-induced ferroptosis ^107^. ROS is equally mitigated by the formation of neutral LDs ^108^ that sequester FFAs and PUFA-OOHs ^72^. Dysregulation of these mechanisms for redox homeostasis triggers pro-apoptotic states ^109,110^ due to damage in the plasma membrane and PS externalization ^66^, a potential activating surface for FXII zymogen (Figure 4C). Indeed, it was reported that PS-induced coagulation of apoptotic cells aids CTCs survival through the activation of coagulation via tissue factor ^111^. With our model, we report a novel organotropic mechanism that with similar metabolic perturbations allows liver metastatic cells to hijack the coagulation system via FXII activation.

Tumor-bearing mice with liver metastasis often presented abdominal distention and bloody ascites that correlated with the elevated expression of FXII (Figure 5G). Malignant ascites has been reported as a symptom of several types of cancer including breast cancer in humans ^112,113^. Ascites could appear because of hepatic hypertension and is frequently associated with portal vein obstruction and deep vein thrombosis caused by liver diseases including liver malignancies ^114–116^. This could explain the pathology observed in E2F5 CKO tumor-bearing mice that present liver metastasis invading the hepatic vasculature and the caudate liver lobe (Figure 6A and 6H). The presence of ascites is common in epithelial ovarian cancer patients (EOCs) ^117^, disease characterized by chronic inflammation ^118^ and with 70% incidence of peritoneal metastasis ^119^. Interestingly, F12 is usually upregulated in the peritoneum of patients with EOC ^118^.

FXII-exposed tumor-associated macrophages are abundant in the peritoneal tissue and favor the invasive capacities and phagocytic potential of EOC ^120^. Notably, our liver metastatic tumors exhibited similar overexpression of F12, elevated infiltration of macrophages (Figure 5H and Supplemental figure 10B), and enrichment of innate immune and pro-inflammatory signatures. Among the enriched signatures observed, cox-macrophages participate in the synthesis of prostaglandins, activated in chronic inflammatory responses ^121,122^; and IL-1β axis, a secreting cytokine that could be induced by FXII ^39^. This pro-inflammatory state is linked to high concentrations of ROS and lipid peroxides in breast cancer ^123,124^ and IL-1β is reported as stimulator of ROS-SRC-MAPK-AP-1 signaling axis ^125^.

In our proposed model, E2F5 transcription factor binds to the promoter of several genes associated with the regulation of lipid metabolism, redox homeostasis, and coagulation. Liver metastatic cells undergo important metabolic adaptations upon interaction with the liver microenvironment. These adaptations generate a high oxidative state characterized by the dependency on external sources of lipids and the inefficient formation of lipid droplets, probably due to the absence of E2F5 global control. This metabolic switch generates abundant lipid peroxides and induces a pro-apoptotic state that increases the exposure of phosphatidylserine in the outer leaflet of the plasma membrane. The elevated stress additionally triggers the production of polyphosphates. E2F5 Knockout also induces the overexpression of the coagulation factor FXII. The exalted concentrations of negative charges due to phosphatidylserine and polyphosphates activate the tumor-derived and the endogenous FXII zymogens, inducing the activation of the intrinsic coagulation cascade. This tumor-associated activation of coagulation confers a protective shield that improves tumor cell migration, immune evasion, facilitates vascular lodging, and liver invasion (Figure 7).

**Figure 7.**
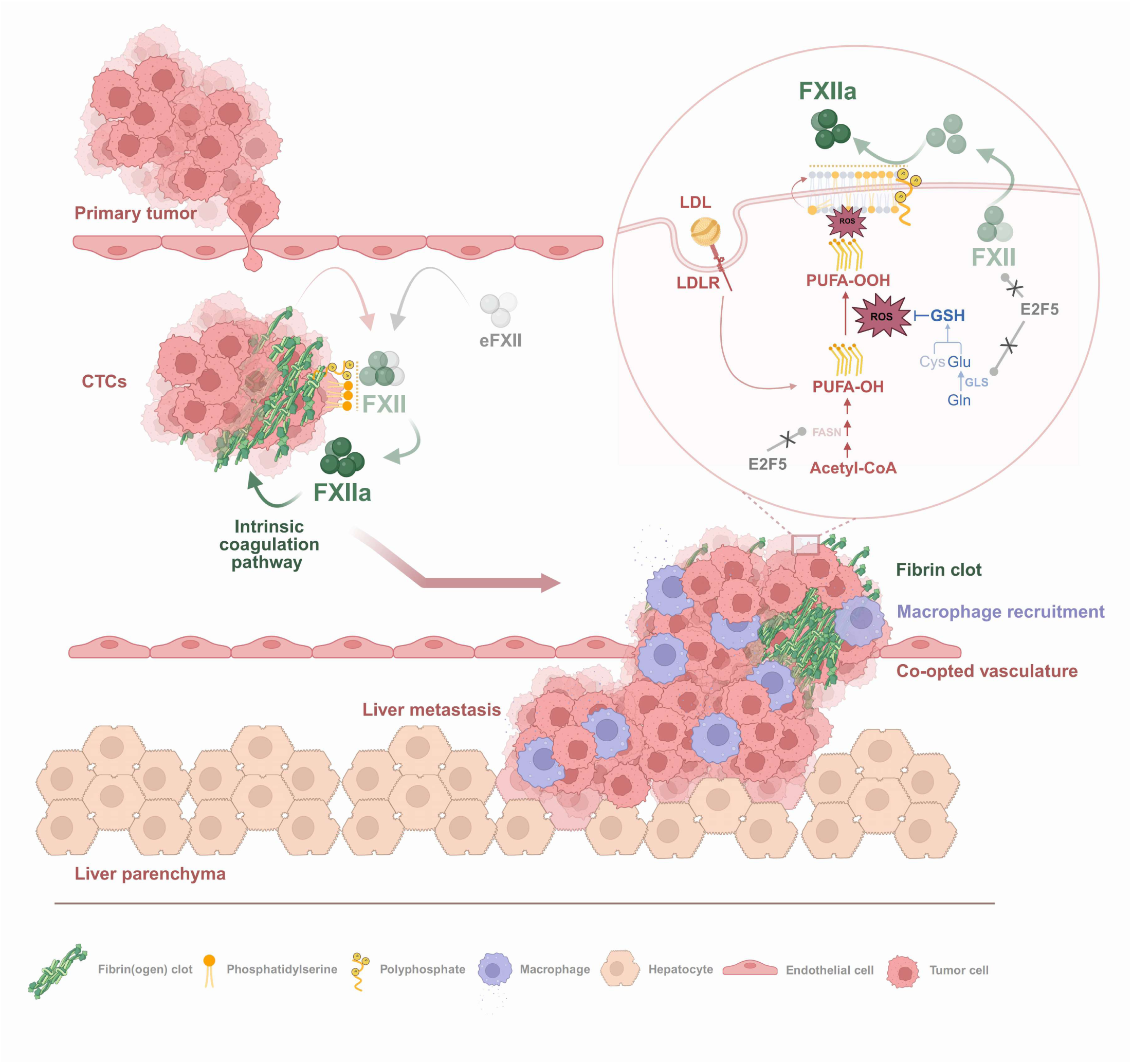
Proposed mechanism of coagulation activation in liver metastasis is globally regulated by E2F5. Breast cancer metastasizes the liver through a coagulation-dependent and E2F5 regulated mechanism. E2F5 transcriptionally regulates F12, and genes controlling lipid and glutamine metabolism. With E2F5 knockout, liver metastatic cells undergo metabolic reprogramming characterized by increased reliance on lipid uptake and synthesis with an impaired formation of lipid droplets. This aberrant state elevates oxidative stress and lipid peroxidation, promoting phosphatidylserine externalization and polyphosphate production. Simultaneously, FXII is overexpressed in the absence of E2F5 and the accumulation of negative charges molecules on cell surface drive FXII activation and initiation of the intrinsic coagulation cascade. As a result, tumor cells drive a procoagulant phenotype that enables migration, immune evasion, vascular retention, and liver metastasis (LDL: Low density lipoproteins; LDLR: LDL receptor; PUFA: Polyunsaturated fatty acid; GSH: Glutathione; Cys: Cysteine; Glu: Glutamate; Gln: Glutamine; FASN: Fatty acid synthase; GLS: Glutaminase; CTCs: Circulating tumor cells; eFXII: Endogenous FXII zymogen; ROS: Reactive oxygen species)

While this work provides new insight into the interaction of breast cancer liver metastasis and coagulation, several aspects require further studies. First, the temporal origin of the liver-tropic phenotype remains to be defined, including whether the pro-coagulant phenotype is selected from pre-existing primary tumor subpopulations, develops during tumor growth in perivascular regions, is induced by the hepatic microenvironment, or is progressively acquired during serial metastatic colonization. Second, while E2F5 loss is linked to F12 overexpression and metabolic remodeling, the complete transcriptional and epigenetic program, including the transcriptional co-factors connecting E2F5 deficiency to these downstream phenotypes remains incomplete. Finally, although heparin treatment supported the relevance of coagulation-associated mechanisms, future studies that test selective anti- FXII or FXIIa drugs are required to determine whether the contact activation represents a direct therapeutic venue for liver metastasis patients as long term heparin therapy presents a bleeding risk for these patients.

## Authors’ contributions

J.G.L. and E.A. conceptualized the study. J.G.L. designed the experimental approach and methodology, performed experiments, analyzed data, generated figures, and wrote the original draft. J.L., B.C., M.J.F., J.R.J., N.T., W.H., and L.Y. contributed to methodology development, technical guidance, and/or experimental design. J.R.J., N.T., J.V., M.M.O., A.J.S., M.A., D.P., D.H., M.Q., S.M., B.T., C.W., and T.K.T. performed experiments. N.T., J.V., M.M.O., and A.J.S. contributed to formal data analysis. J.V. and N.T. contributed to visualization. N.T., M.M.O., B.C., and J.L. contributed to manuscript review and editing. E.A. contributed to experimental design, methodology, manuscript drafting and revision, supervision, and funding acquisition.

## Data Availability

The RNA-sequencing data have been deposited at GEO with accession number GSE330474. CUT&RUN results were deposited at GEO with accession numbers GSE333527 and GSE333695. For the EHR analysis, primary data cannot be shared due to ethical/privacy constraints. Researchers wishing to gain access to individual patient data, may enter into an agreement with TRUVETA. The data used in this study is available to all Truveta subscribers and may be accessed at studio.truveta.com.

## Code availability

All scripts and small auxiliary files required for computational analyses are available at GitHub Jesus-GarciaLerena/E2f5-liver-metastasis-rnaseq-analysis and they will be archived on Zenodo.

## Acknowledgements

We thank all past and present members of Andrechek lab for their discussions and help on this project; Dr. Karl Olson and Dr. Alfred Robison for critical reading of the manuscript; Dr. Sophia Lunt and Lunt lab members Tessa Jordan and Amir Roshanzadeh for providing lab equipment, reagents and consulting; Dr. Kimberly Malloy for participating as Susan Komen patient advocate in this project. We are grateful to Amy Porter and the Laboratory for Investigative Histopathology for the histology services; Confocal microscopy supervisor Melinda Frame of the Center for advanced microscopy for helping with image acquisition; Director of Flow Cytometry core Soo Hyun Ahn for helping with data acquisition; Director of the Quantitative Bio Element Analysis and Mapping (QBEAM) Center Keith MacRenaris, for helping with data acquisition and providing protocols. Illustrations were created in part with BioRender.com.

