## Supplemental Data Legends for "Tumor-derived FXII engages the intrinsic coagulation cascade to support breast cancer liver metastasis"

**Supplemental Figure 1. Histological characterization and transcriptome profile analysis of E2F5 CKO tumor samples.**

(A) Ki67 IHC on enriched, non-enriched tumors, and normal tissue controls (Left panel, scale bars: 100 µm). Proliferative index (Right panel) was calculated on each sample using Image G, and statistical analysis was performed using one-way ANOVA with Tukey post hoc, error bars: Mean with SEM.

(B) CK19 IHC showing tumor cells expressing epithelial markers in lymph node tropic tumors, whereas liver tropic tumors lack detectable CK19 signal under the same conditions (Left panel, scale bars: 100 µm). E-cadherin qPCR analysis on primary cell lines derived from E2F5 CKO tumors (Right panel). Results display three independent experiments performed for each treatment and analyzed by one-way ANOVA with Tukey test post hoc. (Error bars: SD).

(C) Unsupervised clustering of liver metastasis and primary tumor pairs grouped by generation of transplantation and compared to normal liver tissue.

(D) Violin plot showing whole transcriptome alignment distribution of tumor samples and normal liver tissue correlated to predefined hepatocyte identity module (Supplemental Table 1). Overall differences among samples were determined using Kruskal-Wallis’ test, and pairwise comparisons against liver metastatic group were performed using Wilcoxon rank-sum test with Benjamin-Hochberg correction for multiple comparisons.

(E) Top correlated genes on liver metastatic samples encoding hepatocytes-derived plasma proteins.

(F) ssGSEA for cell type signature enrichment on tumor samples and normal liver tissue.

**Supplemental Figure 2. Enriched pathways selection on E2F5 CKO tumors**.

(A) Identification of 33 Lymph node enriched pathways by filtering multiple GSEA comparisons between E2F5 CKO tumors.

(B) Identification of 114 unique liver enriched pathways by filtering multiple GSEA comparisons between E2F5 CKO.

**Supplemental Figure 3. Coagulation genes are upregulated in liver metastasis**.

(A) Heatmap showing top upregulated genes in liver metastatic samples compared to the rest of E2F5 CKO tumors, cell lines, and normal tissue samples.

(B) F3 normalized expression across samples (Error bars: SD).

(C) F12 normalized expression across samples (Error bars: SD).

(D) Several coagulation factors (Pink circles) showing positive expression correlation with F12 in liver metastatic tumors (Pearson correlation).

**Supplemental Figure 4. F12 correlations and E2F5 occupancy on F12 transcript**.

(A) Coagulation factors positively correlating with F12 expresion on lymph node metastasis, primary tumor liver, and non-enriched original tumors (Pink circles, Pearson correlation).

(B) E2F5 CUT&RUN occupancy on F12 transcript located on a reported weak promoter region (Ernst & Kellis, 2012; Gorkin et al., 2020; Perez et al., 2025; Sloan et al., 2016).

**Supplemental Figure 5. Enriched pathways on human breast cancer liver metastasis**.

(A) Top 40 GO terms identified in a distinct human breast cancer liver metastatic cluster using the AURORA US Metastasis Project data (Garcia-Recio et al., 2022).

(B) GSEA showing enriched oncogenic signature on E2F5 CKO liver metastatic tumors.

(C) GSEA showing enriched oncogenic signature on human breast cancer liver metastasis.

(D) Common enriched pathways from C2 curated and C6 oncogenic MSigDB signature gene sets in mice and human liver metastases.

**Supplemental Figure 6. Coagulation genes as part of the GSEA leading edge andPAM50 classification**

(A) Top upregulated genes on E2F5 liver metastasis that are part of the GSEA leading-edge. Coagulation-related genes are colored.

(B) Volcano plot displaying E2F5 CKO liver metastasis leading-edge overexpressed genes equally upregulated in human breast cancer liver metastasis.

(C) PAM50-based molecular subtype classification of human breast cancer primary tumor and Lymph node, lung and liver metastatic pairs. Heatmap showing samples with unsupervised hierarchical clustering of genes and samples. Columns represent individual samples, and rows genes. Top annotations indicate predicted PAM50 molecular subtype and tumor origin.

(D) PAM50-based molecular subtype classification of E2F5 CKO primary, metastatic tumors and derived cell lines. Heatmap showing samples with unsupervised hierarchical clustering of genes and samples. Columns represent individual samples, and rows genes. Top annotations indicate predicted PAM50 molecular subtype and tumor origin.

**Supplemental Figure 7. In vitro clotting assay kinetics and tumor fibrinogen subunit expression**.

(A) In vitro coagulation assay kinetics demonstrate time fibers take to form on E2F5 CKO cell lines, and human breast cancer cell lines MCF-7 and MDA-MB-231 (n=3, Scale bars: 100 µm).

(B) Capillary western blot identifying fibrinogen alpha, beta and gamma subunits in E2F5 CKO liver metastasis and primary breast tumors.

**Supplemental Figure 8. Coagulation activity and FXII expression in E2F5 CKO tumors**.

(A) Fibrin(ogen) density quantification of in vitro clotting assays performed on FXII-deficient plasma. Statistical analysis was performed using one-way ANOVA (p≤0.0001=***, error bars: Mean with SEM).

(B) Fibrin(ogen) density quantification on liver metastatic cells treated with FXII inhibitor CTI. Statistical analysis was performed using one-way ANOVA (p≤0.001=**, p≤0.0004=***, error bars: Mean with SEM).

(C) Representative images of the effect of TFPI on fibrin(ogen) fibers formation in vitro on liver metastatic cells (n=3, scale bar: 100 µm).

(D) FXIIa and thrombin western blot quantification. Statistical analysis was performed using one-way ANOVA (Error bars: Mean with SEM ).

(E) Immunoblot demonstrating the activation of FXII and thrombin over time in liver metastasis conditional medium. Non cells controls medium were included.

(F) FXIIa expression in protein lysate on liver and lymph node metastatic cell lines.

(G) FXIIa IHC corroborates high intratumoral expression of FXII in liver tropic tumors and their derived cell lines compared to normal tissue (Scale bars: 50 and 100 µm).

**Supplemental Figure 9. Fatty acid content and lipid processing in E2F5 CKO tumors.**

**(A)** Intracellular free fatty acids (FFA) concentration in primary tumors and metastatic cells. Statistical analysis was performed using one way ANOVA with Tukey post hoc ( ns=not significant, p≤ 0.02=*, p≤0.008=**, error bars: SEM).

(B) PLIN2 IHC on tumor tissue showing lack of LD reservoirs in liver metastasis (Scale bars: 20 µm).

(C) Immunoblot of LDLR and FASN on lymph node and liver metastasis (Left panel). Densitometry of bands normalized to β-actin are represented as dots and bar plots. Statistical analysis was performed with student T test (Right panel, error bars: SEM).

**Supplemental Figure 10. Tumor immune microenvironment exploration.**

(A) ssGSEA score values distribution for immune cell type classification.

(B) H&E and F4/80 IHC on lymph node and liver lineage over generation of transplantation (*= Tumor-liver interface, scale bars: 100 µm).

(C) TNF-α IHC expression on E2F5 CKO liver tropic tumors. Normal mammary and MMTV-Myc tumors were used as control (Scale bars: 100 µm).

(D) In vitro clotting assay kinetics on liver metastatic cells incubated with TNF-α and IL-1β (n=3, Scale bars: 50 µm).

(E) FXIIa and F4/80 IHC expression correlates on liver tropic tumor (Scale bars: 100 µm).

(F) Representative flow cytometry results in E2F5CKO primary tumors and liver metastasis paired samples displaying the expression of CD8 and CD28 among live singlet CD45+ tumor-infiltrating leukocytes (Left panel). Statistical analysis on 3 paired tumors was performed by Welch’s t test (Right panel, error bars: Mean with SEM).

**Supplemental Figure 11. IMC intensity values.**

Representative images of 3D reconstruction plots showing signal intensity per marker/per area on each E2F5 CKO tumor. Liver metastasis includes a 2D intensity map example generated from Iolite output matrices. Each experimental group was tested three times.

**Supplemental Figure 12. Immune cell type classification and quality control.**

(A-D) 2D generated maps were cell segmented by applying a pseudo-cell grid with tissue coverage established by Cell-ID DNA Intercalator-Ir signal. Raw intensity values were asinh-transformed and assigned to each individual cell for cell type classification and quantification. Image shows representative histograms (Left panels) and correlation heatmaps (Right panels) on representative tumors per group.

**Supplemental Figure 13. In vivo treatment on liver tropic cells.**

(A) Number of CTC colonies on females transplanted on mammary 2/3 or 4 with liver metastatic tumors (Error bars: Mean with SEM).

(B) Primary tumor growth on enoxaparin treated and non-treated females monitored over time and measured in cm are demonstrated. Statistical analysis was performed by two-way ANOVA*,* p≤0.05=*, p≤0.01=**, p≤0.001=***, error bars: SEM).

(C) Correlation between primary tumor volume and number of CTC colonies on vehicle groups. Statistical analysis was conducted with Pearson correlation.

(D) Correlation between Liver metastases size and number of CTC colonies on vehicle groups. Statistical analysis was conducted with Pearson correlation.

**Supplemental Table 1. Hepatocyte identity module**. Selected genes characterizing hepatocyte differentiation, urea cycle, ammonia, hepatocyte-enriched enzymes etc., were selected based on cell markers of [PanglaoDB - A Single Cell Sequencing Resource For Gene Expression Data](https://panglaodb.se/index.html).

**Supplemental Table 2. Unique enriched genes and pathways in liver metastasis on different GSEA comparisons. P**athway selection was restricted to those present on the liver metastasis unique cluster in Figure 2C.

**Supplemental Table 3. Over-represented genes on liver metastatic enriched pathways.** Top upregulated genes are highlighted in blue. Individual DESeq comparisons are detailed in additional excel sheets.

**Supplemental Table 4. LA-ICP-TOF-MS system.** Detailed instrument parameters of ICP-MS (Tofwerk S2), time of flight (Tofwerk S2), and laser ablation (ESL Bioimage 266 nm).
