## Supplemental Figures for "Tumor-derived FXII engages the intrinsic coagulation cascade to support breast cancer liver metastasis"

### Supplemental 1

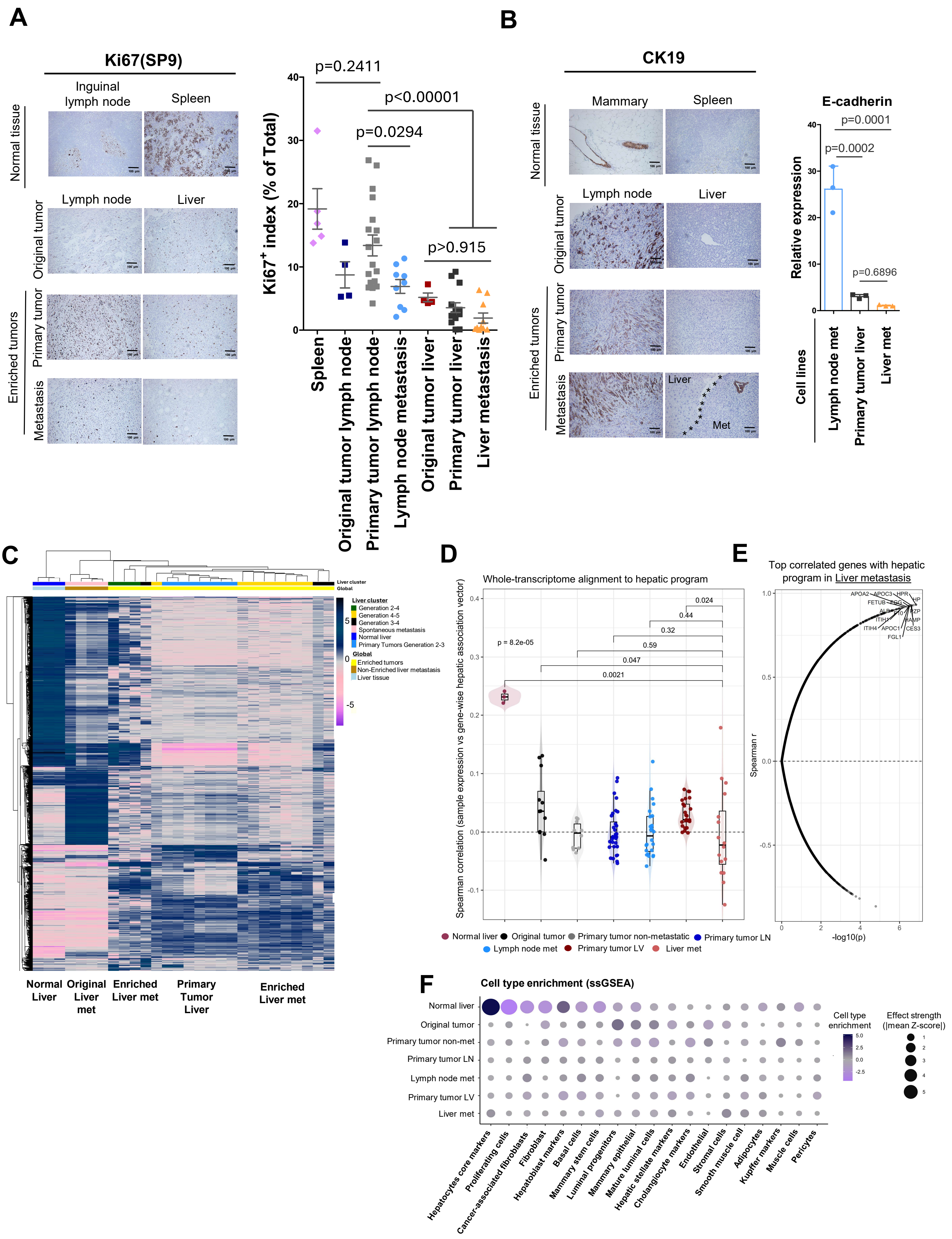

### Supplemental 2

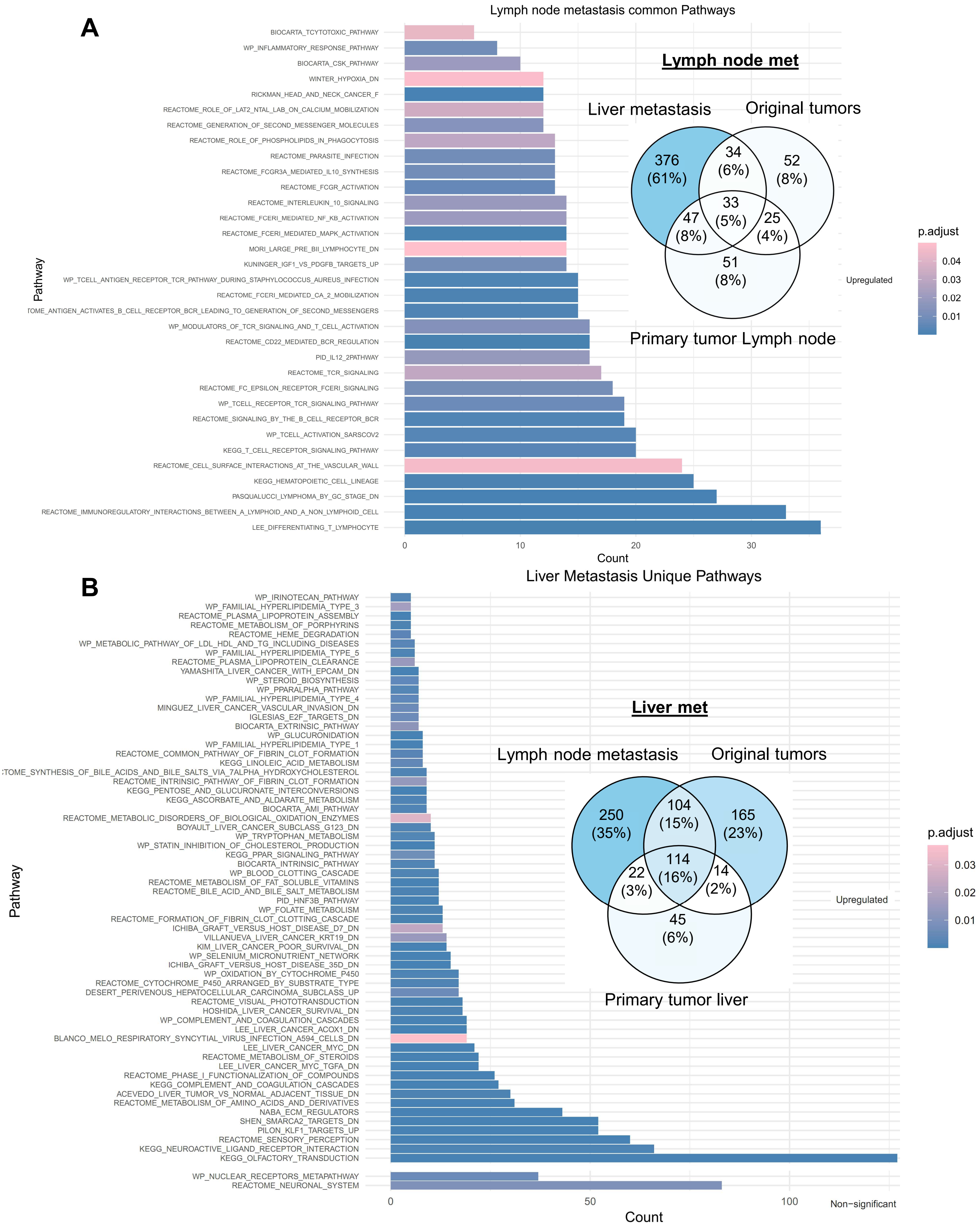

### Supplemental 3

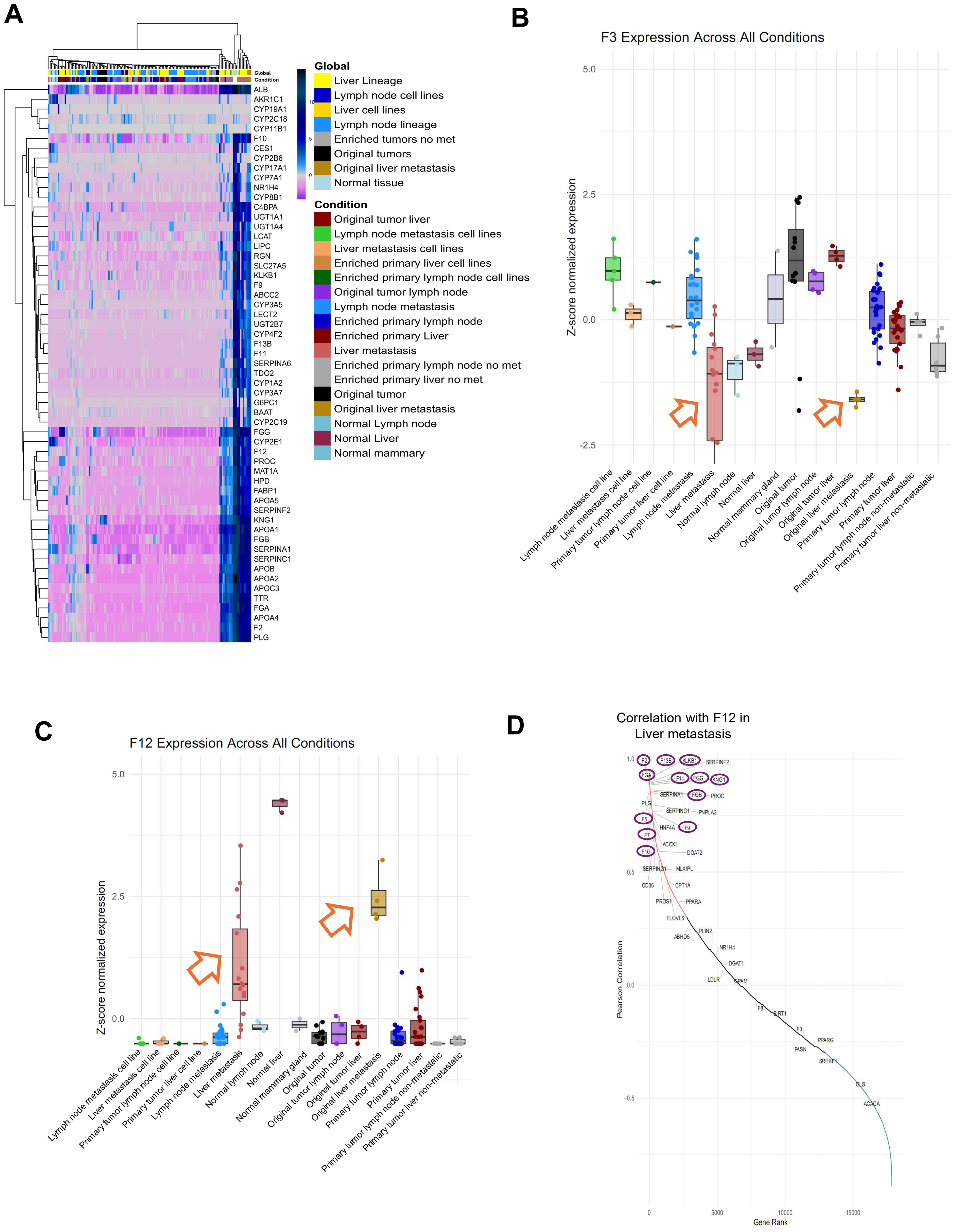

### Supplemental 4

A

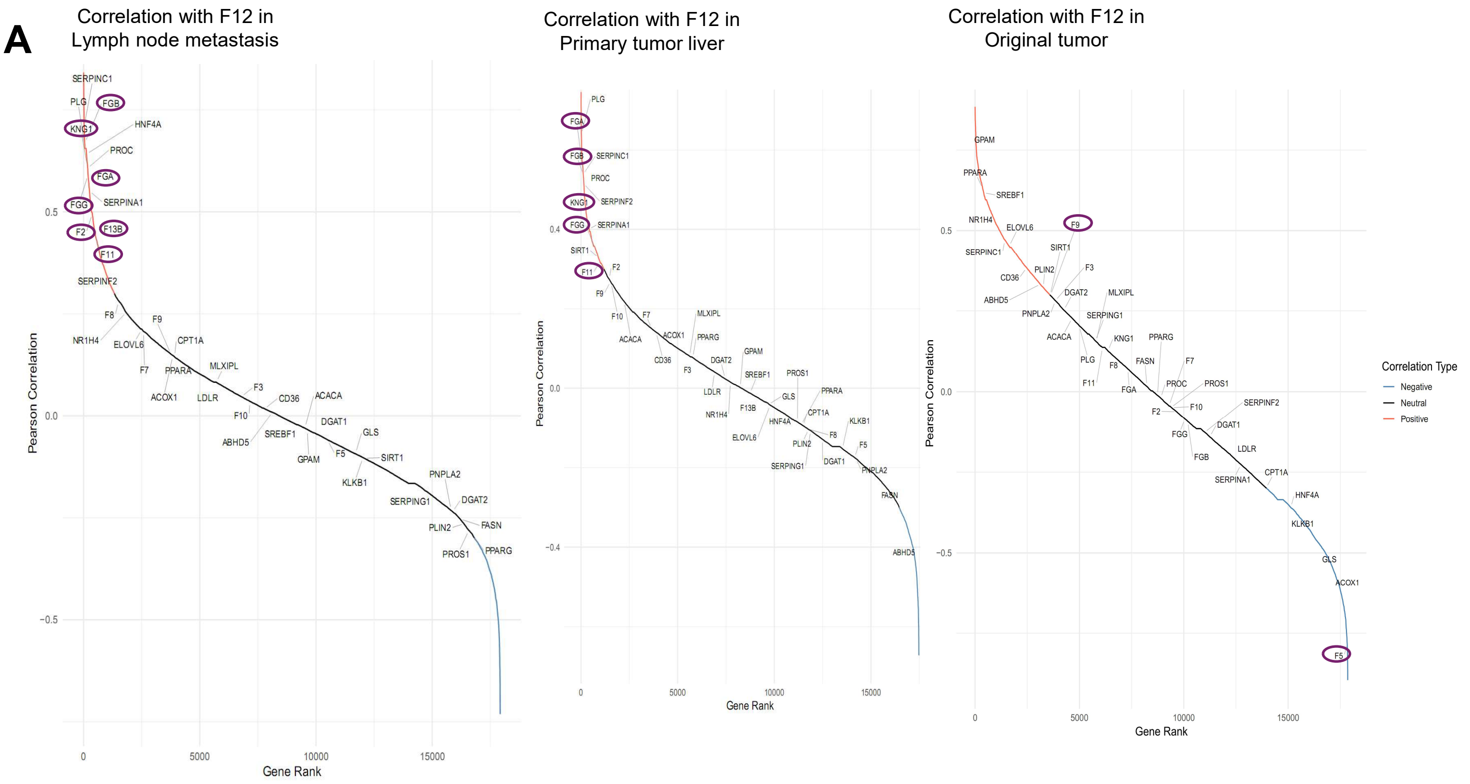


B

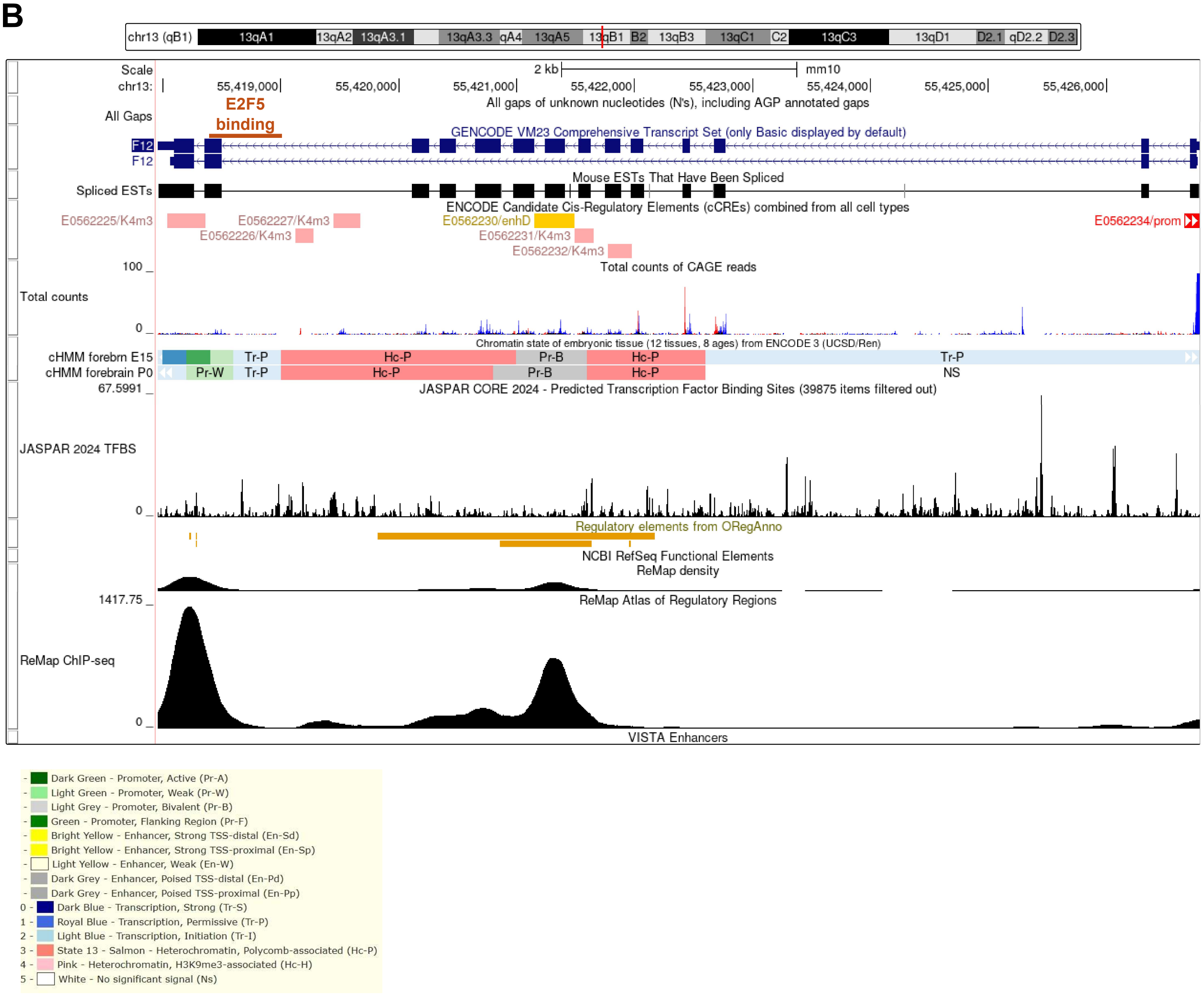

### Supplemental 5

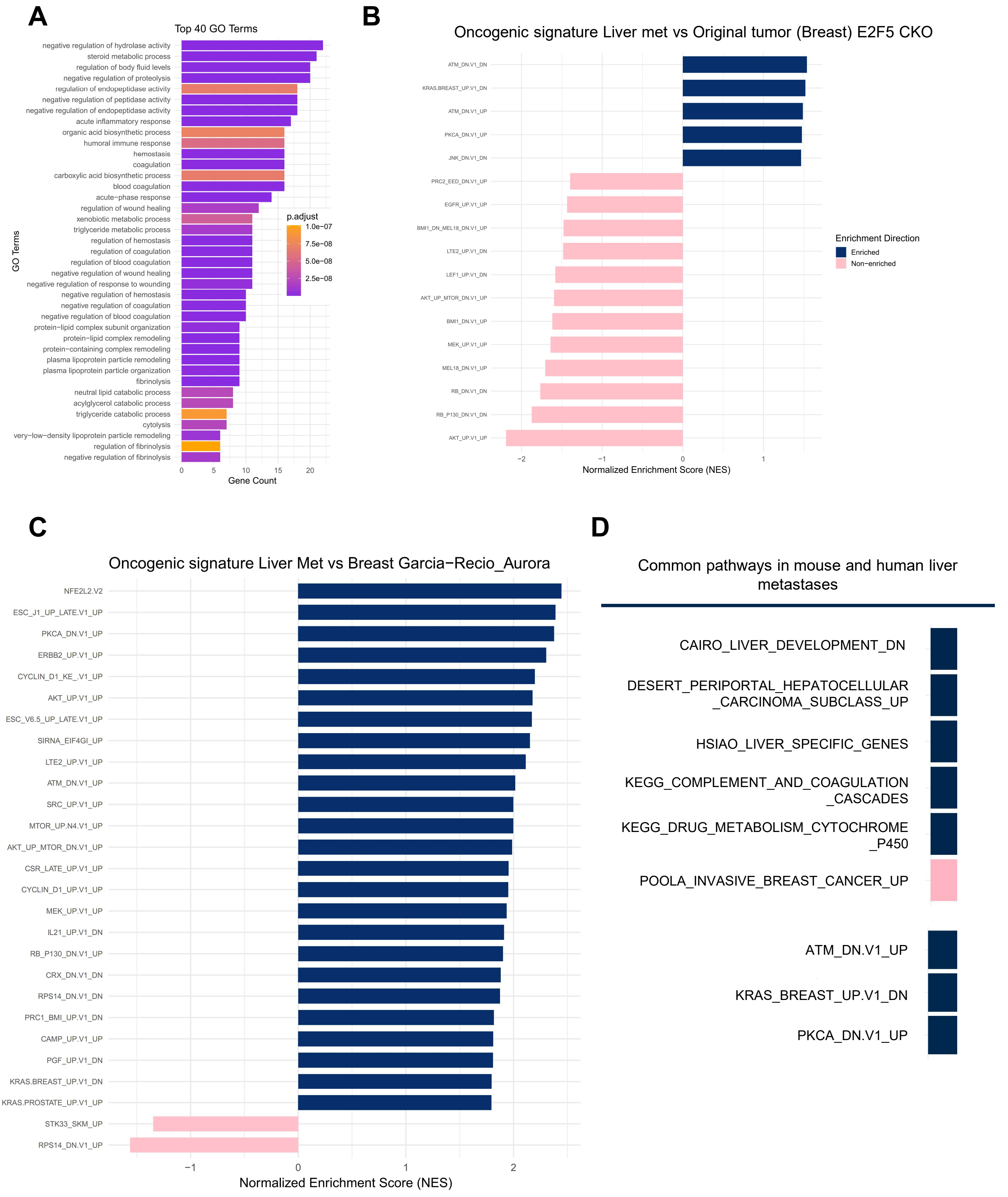

### Supplemental 6

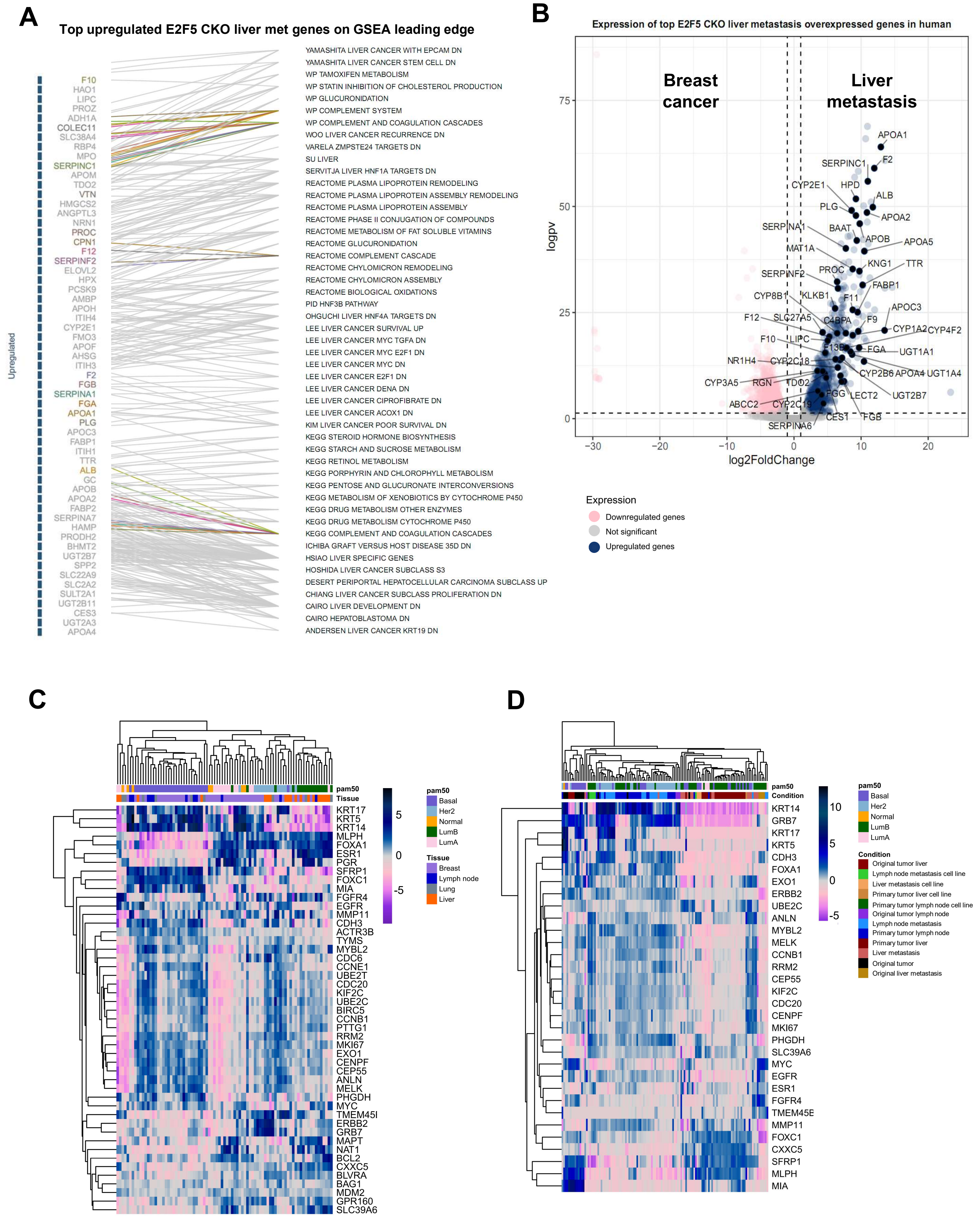

### Supplemental 7

A

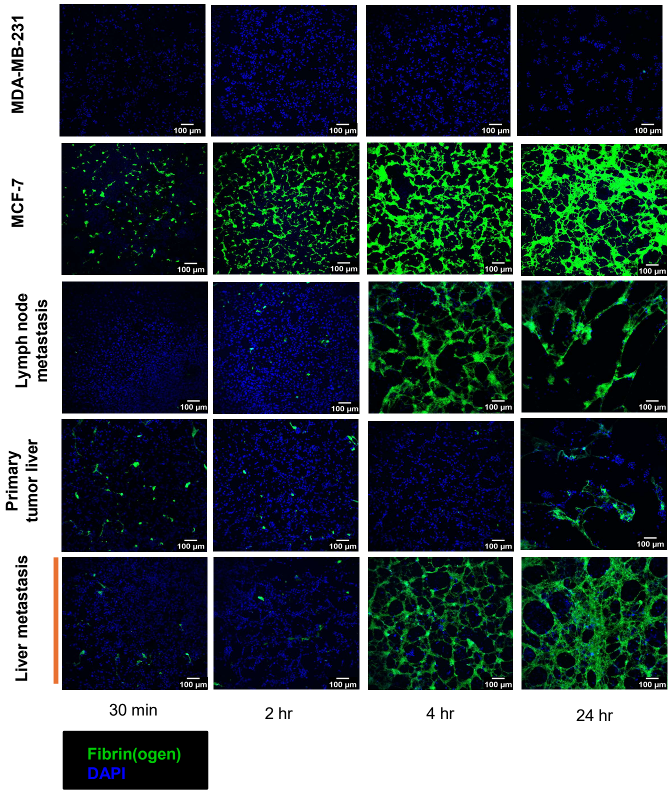


B

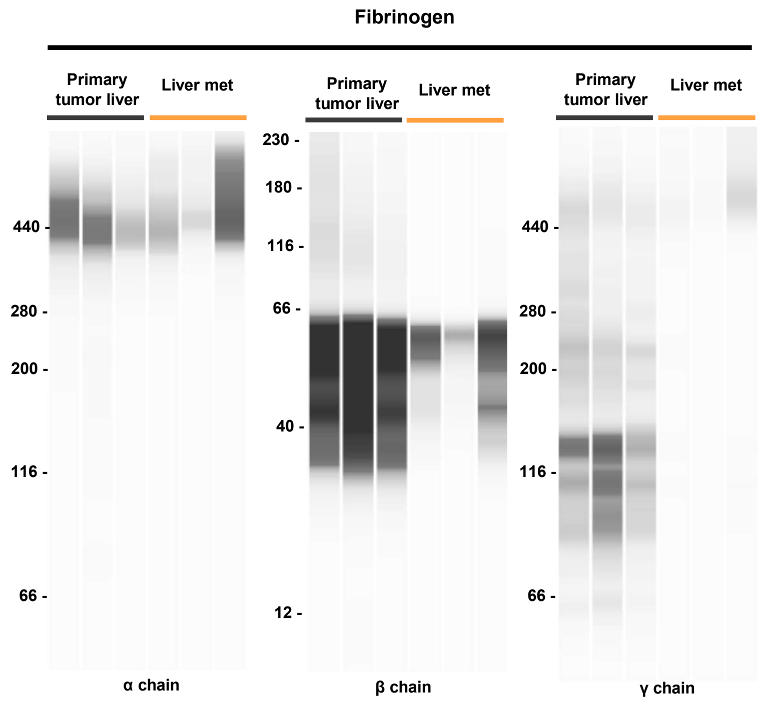

### Supplemental 8

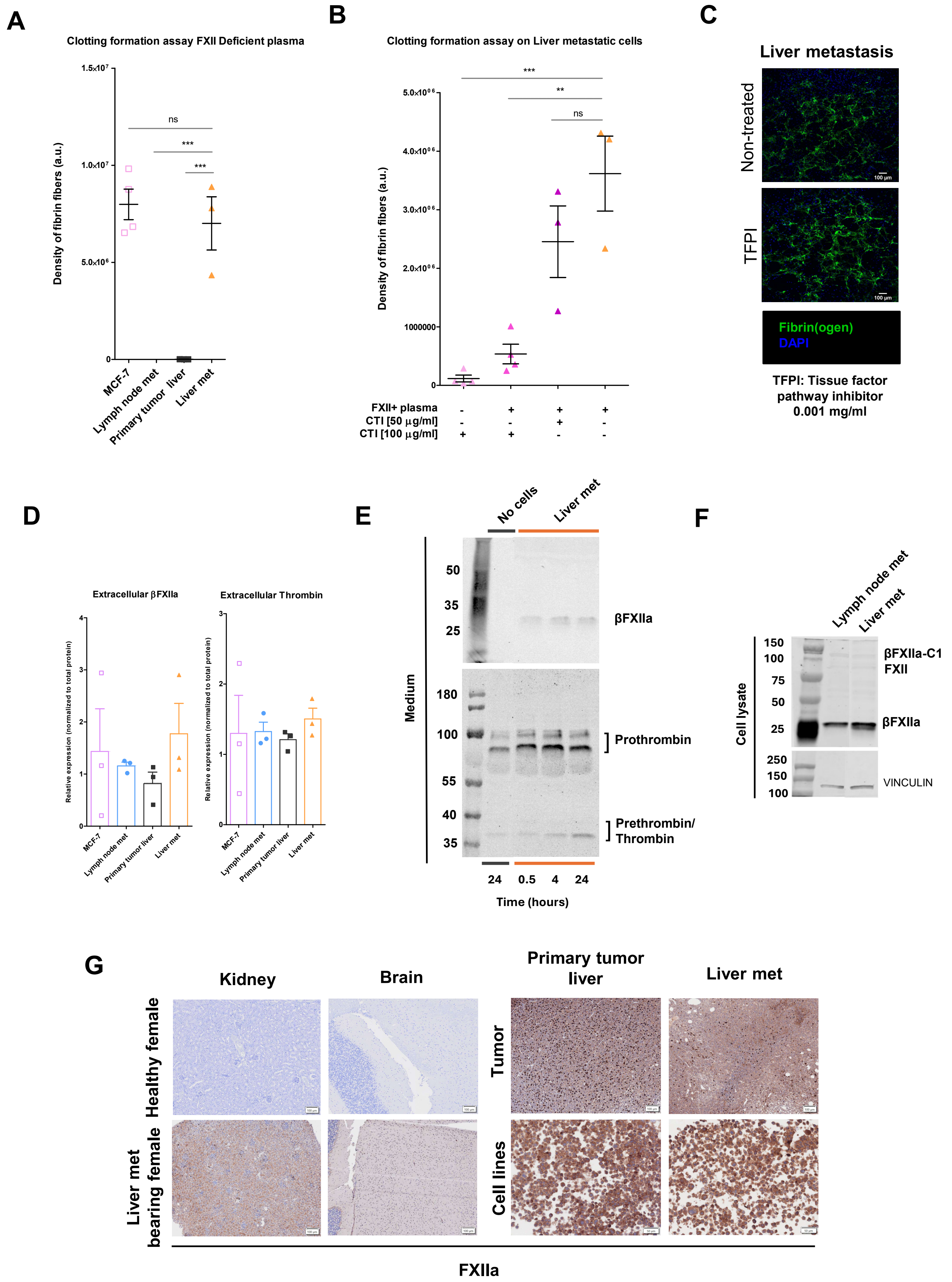

### Supplemental 9

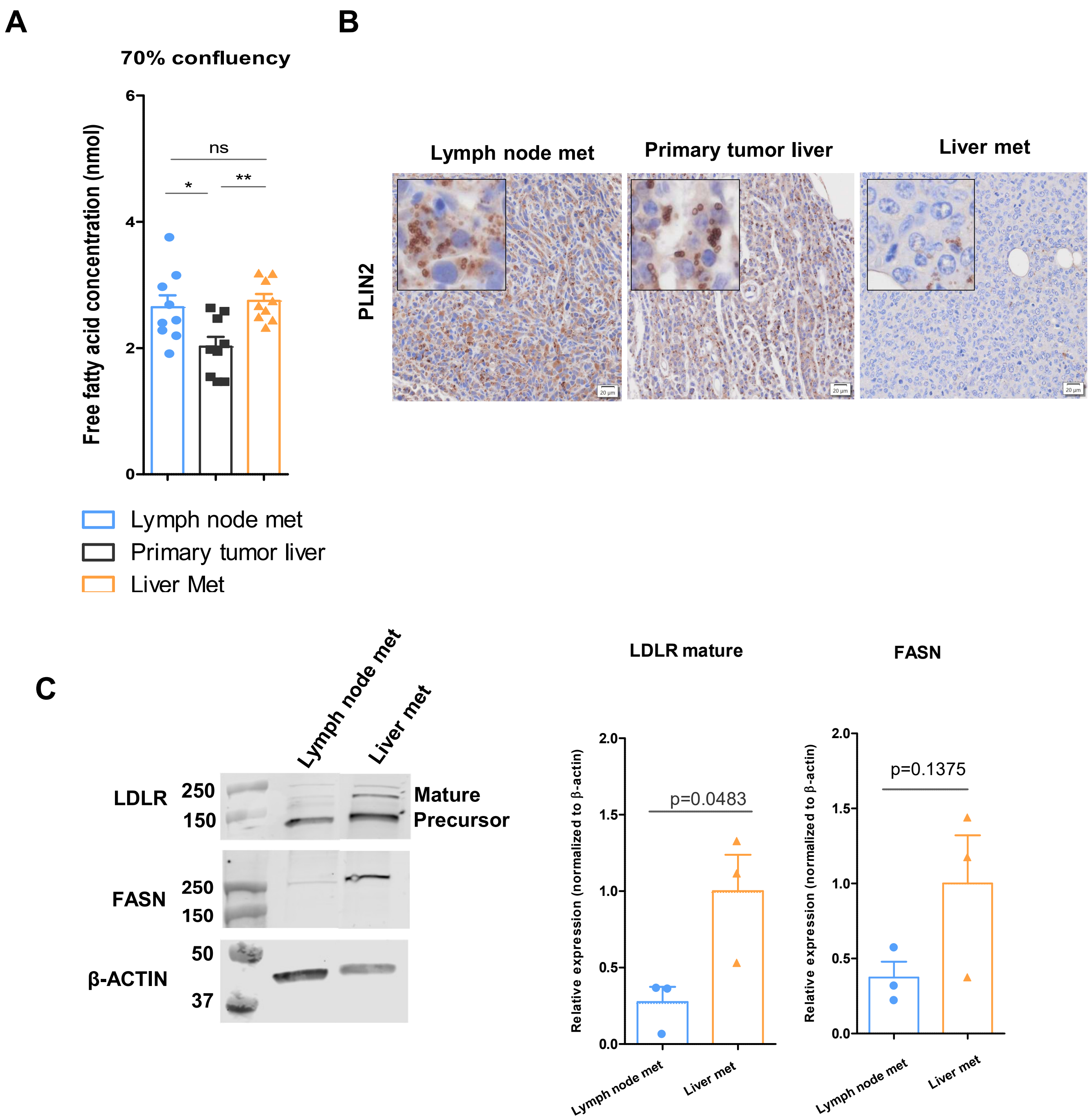

### Supplemental 10

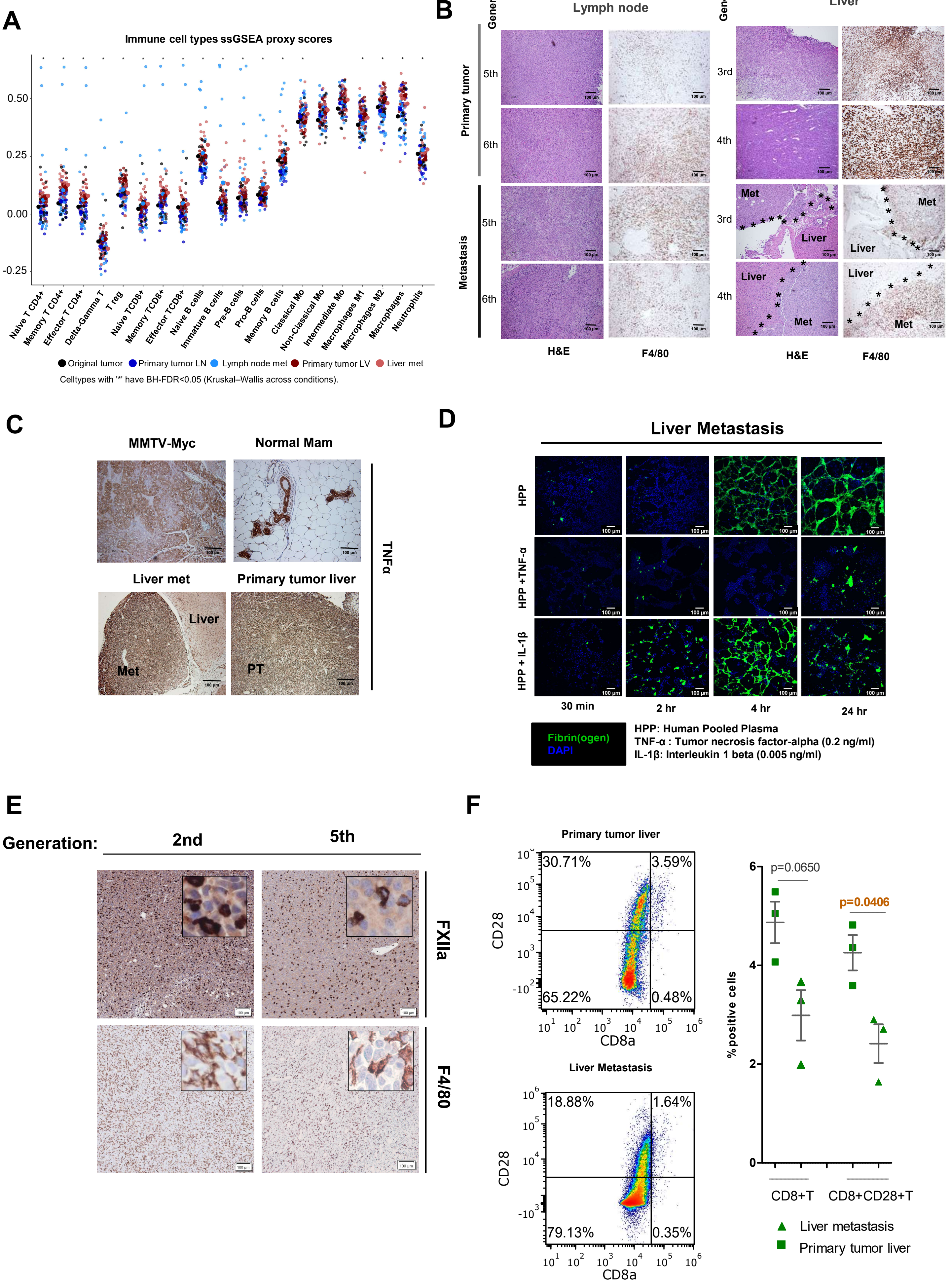

### Supplemental 11

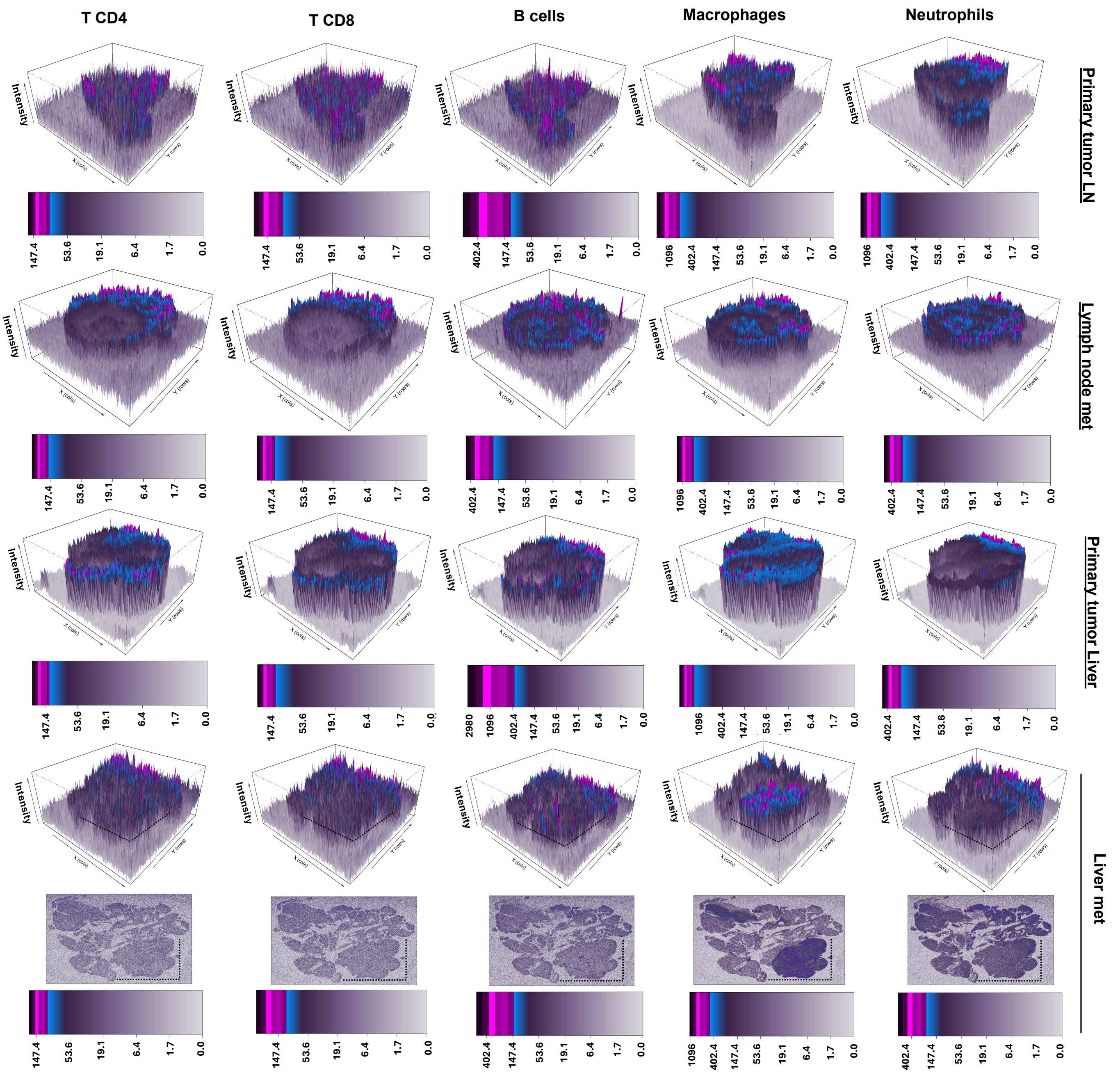

### Supplemental 12

A

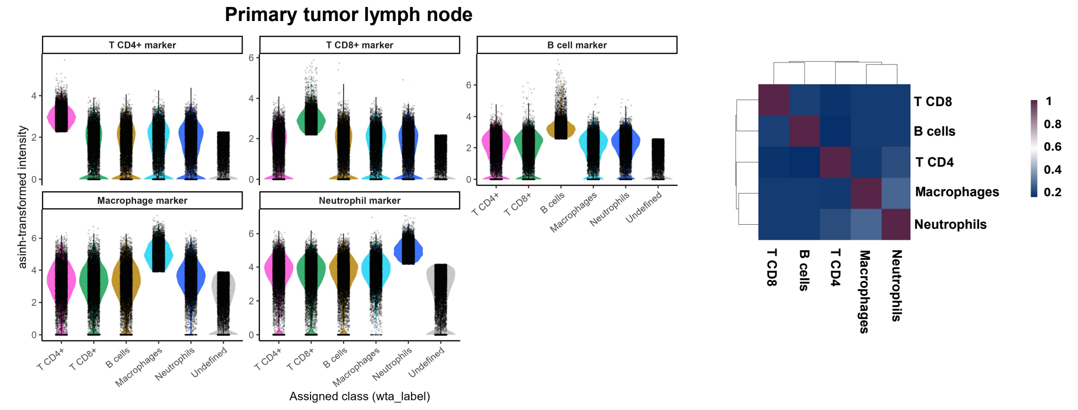


B

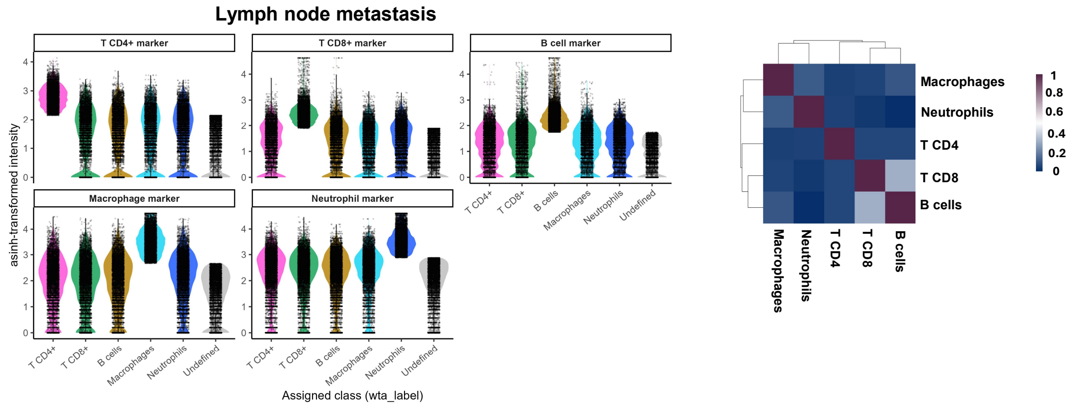


C

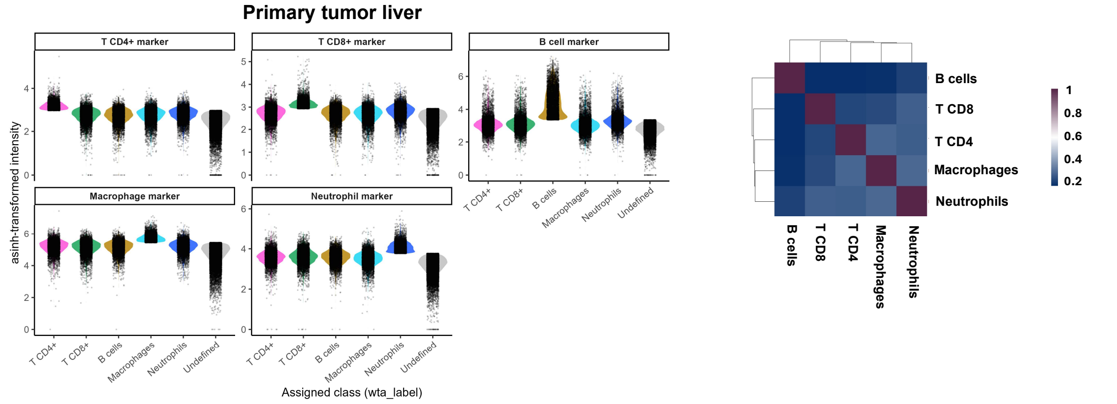


D

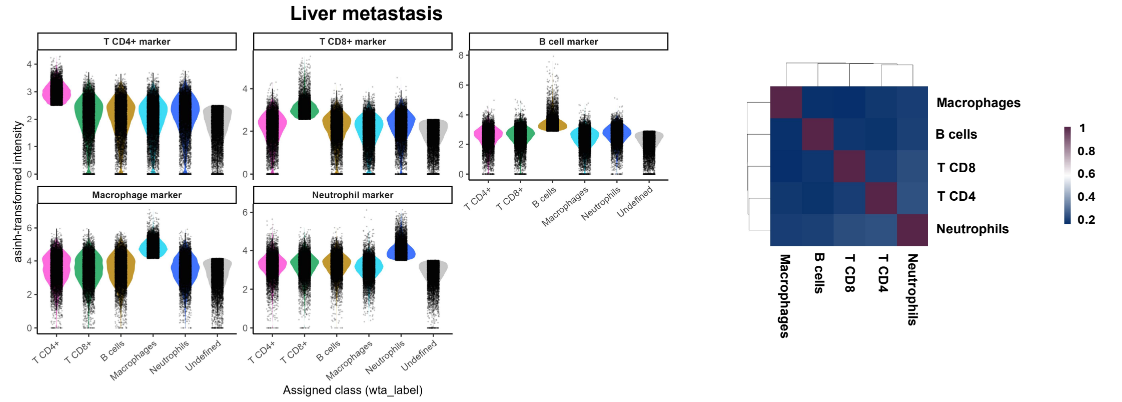

#### Supplemental 13

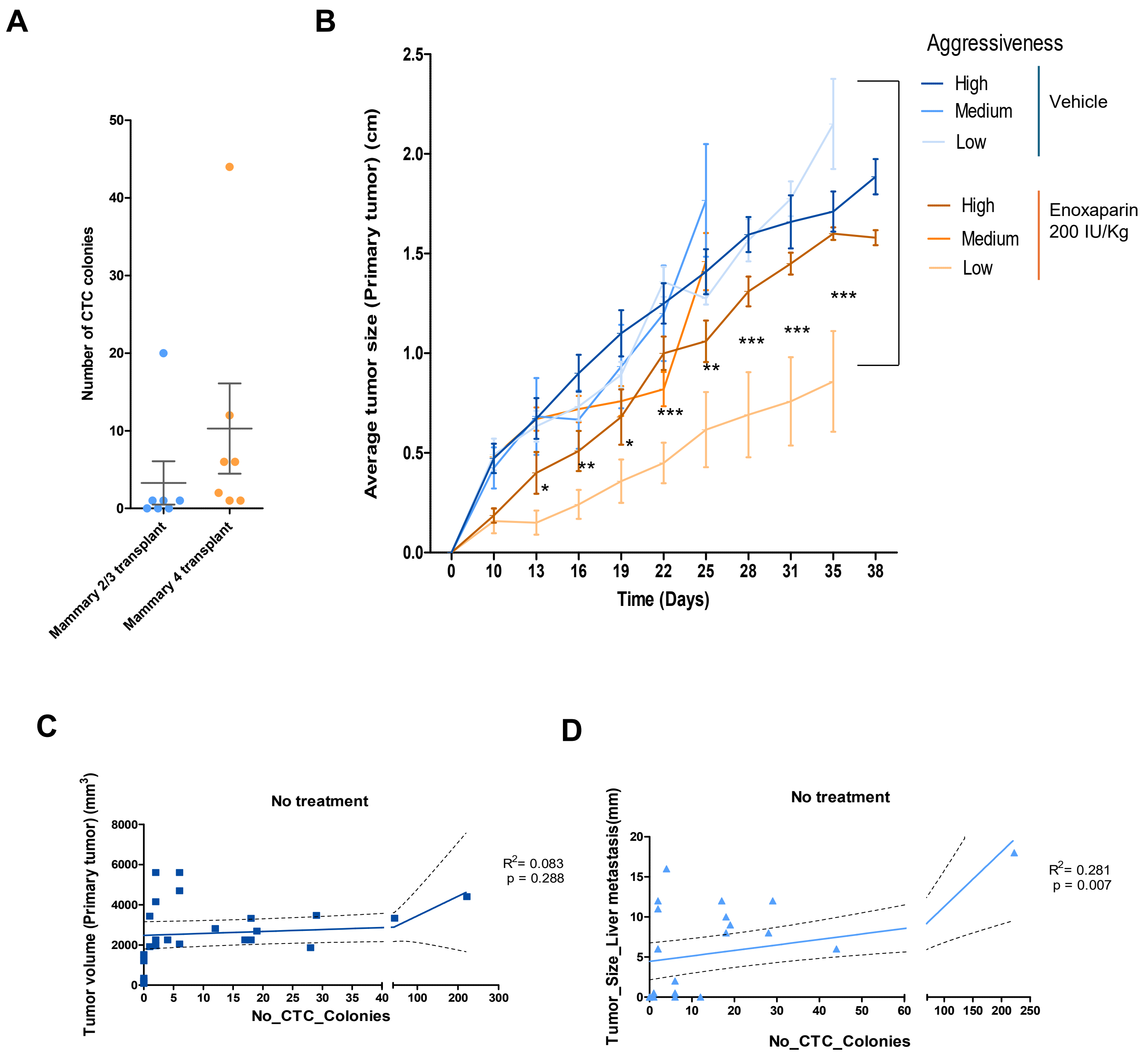
